# The Interplay between Language Development, Short-Term Memory, and Auditory Associative Word Learning in Younger and Older Children

**DOI:** 10.64898/2026.02.15.705078

**Authors:** Samuel H. Cosper, Laura Bachmann, Eleonora Sehmer, Alexandra Steidel, Shu-Chen Li

## Abstract

Auditory associative word learning has been shown in infants and proven to be a difficult task in young adults, where learning is only successful under specific conditions. In order to better understand the transition from successful infant auditory associative word learning to the challenging adult learning, we tested 5-6-year-olds and 9-10-year-olds in a sequential associative task to investigate their ability in associating novel pseudowords with environmental sounds. Additionally, we explored how episodic short-term memory recognition, language development, sex, and musical training affect behavioral and electrophysiological measures of word learning. EEG data were collected to assess word learning in an initial training phase (consistent vs. inconsistent pairings) and a subsequent testing phase (matching vs. violated pairings) with additional button-press reactions for behavioral learning data. While learning effects were seen in the first half of the training phase in younger children, no early effects of learning were found in older children. Only musically trained 9-10-year-olds indicated word learning in the second half of the training phase. In the testing phase, only non-musically trained 9-10-year-olds revealed trend-level N400-like responses. Short-term memory (auditory-verbal, auditory-nonverbal, and visual-nonverbal) and language development improved with age, but only visual-nonverbal short-term memory recognition was positively correlated with improved auditory associative word learning. Unlike cross-modal visual associative word learning, our results, together with earlier findings in infants and young adults, suggest a difficulty in auditory associative word learning beyond infancy, which is sustained from childhood to young adulthood.

## 1 Introduction

In everyday life, humans encounter sensory information from many different modalities including visual, auditory, gustatory, olfactory, and haptic events. Language provides a manner for sharing these experiences with others. Most research on the acquisition of word meaning is conducted largely in cross-modal paradigms with visual objects and spoken lexico-semantic items presented auditorily (e.g., Breitenstein et al., 2005; Friedrich & Friederici, 2008, 2011; Horst & Samuelson, 2008; Junge et al., 2012; McMurray et al., 2012; Sloutsky et al., 2017; Smith & Yu, 2008; Taxitari et al., 2019; Werker et al., 1998). There are also studies, albeit few, conducted in word learning in other modalities such as auditory (e.g., Cosper et al., 2020, 2022, 2024), tactile (e.g., Miller et al., 2018; Schmidt et al., 2019), olfaction (e.g., Vanek et al., 2021), and gustatory (e.g., Yan et al., 2021). Recent research has attempted to investigate if words with auditory references like *thunder* are encoded similarly to words with visual references like *lightning* in both infants (Cosper et al., 2020) and in young adults (Cosper et al., 2022, 2024). The goal of this current study is to further the understanding of the mechanisms behind the mapping of spoken pseudowords onto auditory objects (auditory associative word learning) by attempting to find when children are no longer as sensitive to auditory information as in infancy and behave more like adults in their associative learning patterns (cf. Cosper et al., 2022). To pursue this goal, we investigated auditory associative word learning, modality specific short-term memory recognition, and language development in younger (5-6-year-olds) and older (9-10-year-olds) children.

For word meaning acquisition to occur, a relationship between a *signifié* (signified; e.g., an object) and a *signifiant* (signifier; e.g., a label), be it a spoken word, written word, or signed word, must be established (cf. McMurray et al., 2012; Quine, 1960; Samuelson & McMurray, 2017). For this, various learning models have been proposed, including association-based and inference-based models (cf. Sloutsky et al., 2017). Their relative dominance may change across the lifespan, with infants relying more on associative learning processes (McMurray et al., 2012; Sloutsky et al., 2017; Smith & Yu, 2008; Werker et al., 1998), and adults relying on inference-based learning, such as hypothesis testing (Xu & Tenenbaum, 2007). However, this distinction might not be clear-cut, as evidence further suggested other mechanisms may be at work (Waxman & Gelman, 2009). Further studies indicated that associative mechanisms play an important role not only in childhood (e.g., Bergelson & Swingley, 2012; Bortfeld et al., 2005; Johnson, 2016; Johnson et al., 2014; McMurray et al., 2012; Shukla et al., 2011; Sloutsky et al., 2017; Tincoff & Jusczyk, 1999, 2012) but also in adulthood (e.g., Regier, 2005; Sloutsky et al., 2017).

Although behavioral measures can be used to measure word learning, neuroimaging measures can also provide further important insight, specifically event-related potentials (ERPs) derived from electroencephalography (EEG). ERPs may be sensitive to aspects of word learning, e.g., through reflecting lexical knowledge, that might be overlooked by behavioral measures (McLaughlin et al., 2004). In particular, the N400 component marks a violation of expectation. It is described as a negative-going waveform which typically peaks around 400 ms after the onset of the target stimulus and with a centro-parietal electrode site distribution (Kutas & Hillyard, 1980) and can be found both during training (Mestres-Misse et al., 2007) and after learning (for reviews, see Junge et al., 2021; Kutas & Federmeier, 2000, 2011). Functionally, N400 models have been found to be related to probabilistic representation of meaning (Rabovsky et al., 2018), processing of related predictions and prediction error (Bornkessel-Schlesewsky & Schlesewsky, 2019), predictability (semantic-level), plausibility (sentence-level), and similarity (low-level semantic relationship, based on co-occurrence; Nieuwland et al., 2020). The latency of the N400 in adults has also been suggested to be influenced by factors such as object modality (Bechtold et al., 2023; Cosper et al., 2022), tension and contradiction in personal semantics (identified by the authors as the N500; Schöne et al., 2018), as well as the use of frontal, but not lateral, gestures (He et al., 2020). Furthermore, in infants factors such as age, stimuli-specific processing demands, and vocabulary have been identified as being influential to the latency, amplitude, and topological distribution of an N400-like effect (Junge et al., 2021). As such, the N400 component is a suitable component to measure word meaning acquisition both during and after associative word learning paradigms.

Recent research into auditory associative word learning has revealed a possible spectrum of learning that changes across development. Infants between 10-12 months can not only associate environmental sounds with novel pseudowords, but the topological distribution is much broader than similar studies conducted with novel associations of visual objects and in a somewhat earlier time window (300-400 ms post stimulus; Cosper et al., 2020). Adults, on the other hand, exhibit difficulties in building sound-pseudoword associations and, instead, indicate a preference for visual objects and stronger associations when stimuli are presented simultaneously instead of sequentially (Cosper et al., 2022). When considering these findings as two points on a developmental trajectory, two main questions can arise: (i) when does auditory associative word learning shift from mirroring the successful infant-like learning and begin to resemble adult-like learning and (ii) what are the potential factors contributing to such a shift. Several influential factors have already been identified, including sensory modality and stimulus order (Cosper et al., 2022, 2024). Other factors have been suggested, but not directly tested, including sensory dominance, short-term memory (STM), and STM maintenance (Cosper, 2020; Cosper et al., 2020, 2022, 2024). Further factors to be considered include time spent in formal education (cf. Nava & Pavani, 2013) and vocabulary size (cf. Junge et al., 2021).

In regard to perceptual modality, this study focuses mainly on the auditory modality, with comparisons to visual and audiovisual multimodal perception. With respect to these modalities, there are differences in processing and in learning across development. Although both infants and children are able to apply statistical learning to both auditory and visual stimuli (e.g., Kirkham et al., 2002; Saffran, 2002; Saffran et al., 1996, 1999), there are differences in processing and learning between the modalities. Infant and adult statistical learning has been reported to be superior in the auditory modality compared to the visual-temporal modality, but similar to the visual-spatial modality (Conway & Christiansen, 2005, 2009; Emberson et al., 2011, 2019). Between the visual and auditory modalities, adults are, however, shown to have a stronger object memory for visual objects than auditory objects (Cohen et al., 2009; Yuval-Greenberg & Deouell, 2007, 2009). Differences in processing also occur within modalities. The auditory modality can be divided into auditory-verbal (i.e., language) and auditory-nonverbal (e.g., environmental sounds) information. It has also been shown that adults process auditory-verbal and auditory-nonverbal information differently (Calignano et al., 2021; Lukics & Lukács, 2022; Thiessen, 2010; van Petten & Rheinfelder, 1995). While the early developmental trajectory of statistical learning improves in the auditory modality, it does not improve the visual modality for infants between 8-10 months (Emberson et al., 2019). However, for 5-12-year-olds, statistical learning improves in the visual and auditory-nonverbal modalities, while measures of auditory-verbal statistical learning do not significantly increase (Raviv & Arnon, 2018; Shufaniya & Arnon, 2018).

Furthermore, there are attentional differences during multimodal processing both during early development and in adulthood, which can be seen in sensory dominance effects. In development, children attend more to auditory stimuli than visual, except when visual stimuli are presented more saliently than the auditory stimuli (e.g., Lewkowicz, 1988b, 1988a; Robinson & Sloutsky, 2004, 2019; Sloutsky & Napolitano, 2003). Whereas in adulthood, a general visual dominance has been reported (Hirst et al., 2018; Robinson & Sloutsky, 2004; Sloutsky & Napolitano, 2003; Soemer & Saito, 2015); however, recent research has indicated that the shift to a visual dominance may only occur in older adults, and younger adults might still exhibit an auditory dominance (Barnhart et al., 2018). With consideration to the breadth of research in which young adults were found to exhibit a visual dominance, Nava and Pavani (2013) have suggested that the switch from an auditory dominance to a visual dominance occurs between the ages of 7 and 9, which may also be influenced by time spent in formal education. Although the current study focused on auditory associative word learning and not perception, the influence of sensory dominance may contribute to the shift from infants being able to successfully learn novel labels for auditory events (Cosper et al., 2020) to adults not reflecting similar learning abilities in the same experimental capacity and benefitting from the labeling of visual objects (Cosper et al., 2022, 2024).

Memory may also play a role in the shift from the infant-like associative word learning to the adult-like learning of auditory stimuli (Bechtold et al., 2023; Cosper, 2020; Cosper et al., 2022, 2024). This can be seen in memory processes for auditory events in young and middle-aged adults, which has been shown to be weaker than visual memory (Bigelow & Poremba, 2014; Cohen et al., 2009; Talamini et al., 2022; Yuval-Greenberg & Deouell, 2007, 2009). Even though modality differences in STM may conflate the outcome of statistical learning, modality-matched performance in the visual and the auditory modalities are comparable between children (7-9-years-old) and young adults (Saffran, 2002). Furthermore, processing differences are found between auditory-verbal and auditory-nonverbal information (Calignano et al., 2021; Lukics & Lukács, 2022; Thiessen, 2010; van Petten & Rheinfelder, 1995); whereas adult performance accuracy in verbal STM tasks are associated with higher accuracy than with nonverbal stimuli in both the auditory and visual modalities (Talamini et al., 2022). How can this be? Due to the fleeting nature of the auditory modality, it is vital that STM be maintained. STM maintenance in the auditory modality is generally described in the cognitive mechanism of the phonological loop, a mechanism reliant on vocal or sub-vocal rehearsal and can be applied to both auditory-verbal and auditory-nonverbal stimuli (for a review, see Baddeley, 2012). However, a different cognitive mechanism may be responsible for the maintenance of non-verbal information, namely auditory imagery (for a review, see Hubbard, 2010), which does not necessarily rely on self-rehearsal strategies. The notion of subsystems of auditory STM was empirically investigated and results suggest not only the existence of these distinct subsystems, but their importance for word learning (McQueen et al., 2020). As hypothesized in Cosper and colleagues (2022), the application of both methods of memory maintenance (auditory imagery for auditory-nonverbal stimuli and the phonological loop for auditory-verbal stimuli) may cause interference in STM, contributing as a factor to auditory-sequential associative word learning being more difficult for adults (for further information on stimulus-related factors to word learning including saliency and memorability of auditory-verbal and auditory-nonverbal information, see Cosper et al., 2024). However, it is unknown to what extent this may be true and if children are also affected by this in a similar manner. In addition to the possible influence of these multiple cognitive aspects, the saliency of spoken language (cf. Cosper et al., 2024) may also affect how auditory stimuli are maintained. This, however, is considered in the modality specific STM tasks in current set of experiments.

Taken together, to address the age-related, language-related, and memory-related developmental gaps in the literature, we designed the current study to delineate a clearer picture of the shift from infant-like learning to adult-like learning. Each participant completed five tasks: a developmental language assessment (SET 5-10; Petermann, 2012; Rißling et al., 2013), an EEG auditory associative word learning task (AAWL task), and three behavioral episodic short-term memory recognition tasks (STM tasks): auditory-verbal, auditory-nonverbal, and visual-nonverbal. The 5-6-year-old and 9-10-year-old age groups were chosen because of documented statistical-learning developmental tracks (Raviv & Arnon, 2018; Shufaniya & Arnon, 2018), the shift from auditory dominance towards visual dominance (between 7 and 9 years of age; Nava & Pavani, 2013), and evidence of multisensory integration only occurring after the age of 8 (Gori et al., 2008; Nardini et al., 2008). Our analysis was split into hypotheses and exploratory analyses. We hypothesized that both younger children (5-6-year-olds) and older children (9-10-year-olds) are able to map novel pseudowords onto environmental sounds. We also expected to find differences between the groups in the amplitude and/or latency of the N400 in the testing phase, whereas younger children have larger and earlier ERP effects than older children. Additionally, we attempted to disentangle the factors beyond object modality that might influence word learning in exploratory analyses. First, we explored STM as a factor. Here, we assessed auditory-verbal, auditory-nonverbal, and visual-nonverbal STM recognition in an exploratory manner. Next, we explored possible influences of language development on auditory associative word learning and STM recognition. In order to quantify language development, we conducted a language development test battery (SET 5-10; Petermann, 2012; Rißling et al., 2013). Finally, we explored biological sex (henceforth sex) differences, as behavioral differences have been found in both children and adults (Kaushanskaya et al., 2011, 2013), and differences due to musical training, as musical training affected associative word learning in the visual modality in both children and young adults (Dittinger et al., 2016, 2017) as well as young adults in the auditory modality (Cosper et al., 2024).

## 2 Materials and methods

### 2.1 Participants

Two groups of children participated in the study. The younger group consisted of a total of 57 children (27 males; 30 females) ranging from 5-6 years of age. The older age group included a total of 54 children (29 males; 25 females) ranging between 9-10 years of age. In order to be included in the final dataset, participants had to complete all five tests and retain 50% of each trigger type in the EEG word learning task after preprocessing (including artifact rejection and Independent Component Analysis (ICA); see Table 1 for a list of triggers, their averages, and the percentage of trials retained). A total of 11 participants in the younger group were excluded from the final dataset due to technical reasons (*n* = 2) and for not meeting the minimum requirement of epochs after preprocessing (*n* = 9). The final dataset included 46 younger children (22 males; 24 females) with ages ranging from 5.06 – 6.97 years of age (*M* = 6.19 years; *SD* = 0.56 years). In the younger age group, only 4 children were reported to having learned a musical instrument for at least 1.5 years. As for the group of older children, 7 participants were excluded from the final analyses due to not finishing all five tests (*n* = 6) and due to not meeting the minimum number of epochs per experimental condition after preprocessing of the EEG word learning task (*n* = 1). The final dataset consisted of 47 older children (24 males; 23 females) with ages ranging from 9.01 – 10.52 years (*M* = 9.78 years; *SD* = 0.43 years). In the older age group, 19 participants were reported to learning to play a musical instrument for at least 1.5 years.

**Table 1:** Overview of participants by age group along with sample size, the number of participants musically trained for at least 1.5 years (at the time of testing), and average number of trials per condition included in the final dataset for the EEG word learning task. The average number of trials retained for each condition after preprocessing are given along with the percentage of total number of trials retained. Standard deviation was also indicated for all averages. Musical training was determined by parental questionnaire and only included if training was still ongoing and for consecutive time period of at least 1.5 years.

| Participant Group | Sample Size | Average Age (SD) | Musically Trained Participants | 1<sup>st</sup> Half Training – Consistent: 32 total trials (SD) | 1<sup>st</sup> Half Training – Inconsistent: 32 total trials (SD) | 2<sup>nd</sup> Half Training – Consistent: 32 total trials (SD) | 2<sup>nd</sup> Half Training – Inconsistent: 32 total trials (SD) | Testing – Matching: 16 total trials (SD) | Testing – Violated: 16 total trials (SD) |
| --- | --- | --- | --- | --- | --- | --- | --- | --- | --- |
| 5-6-year-olds | 46 | 6.19 years (0.56 years) | 4 participants | 26.76 trials<br>83.6%<br>(2.97 trials) | 27.48 trials<br>85.9%<br>(2.39 trials) | 24.46 trials<br>76.4%<br>(3.06 trials) | 24.28 trials<br>75.9%<br>(3.35 trials) | 12.13 trials<br>75.8%<br>(2.19 trials) | 11.61 trials<br>72.6%<br>(2.58 trials) |
| 9-10-year-olds | 47 | 9.78 years (0.43 years) | 19 participants | 28.15 trials<br>88%<br>(2.31 trials) | 28.4 trials<br>88.8%<br>(2.45 trials) | 27.17 trials<br>84.9%<br>(3.06 trials) | 27.62 trials<br>86.3%<br>(3.08 trials) | 12.96 trials<br>81%<br>(1.8 trials) | 12.7 trials<br>79.4%<br>(1.88 trials) |

All participants were contacted through the database of the Lifelab at the Technische Universität Dresden, were native speakers of German, right handed (in accordance to the Edinburgh Handedness Inventory, presented in German; Oldfield, 1971), reported normal hearing and normal/corrected eye-sight, and did not report any neurological conditions (ADHD, learning impairments, language impairments, dyslexia, depression, or any other neurological or cognitive diagnoses) or any prescription medication that would affect memory or attention. Furthermore, all participants were blind to the goals of the study before participation; however, the participants and parents were informed of the goals after the completion of the five tasks. Before participation, parents and children gave written informed consent. All participants were compensated for their time (a total of three hours over two visits) with 12 Euro per hour for a total of 36 Euro. The experiment was approved by the Ethics Committee of the Technische Universität Dresden (application number: SR-EK-169042023) and conformed to all aspects of the Declaration of Helsinki (World Medical Association, 2013).

### 2.2 Stimuli

#### Auditory Associative Word Learning (AAWL) Task

Stimuli used in the associative word learning task of the current experiment included 16 environmental sounds (from the NEESTI database; Hocking et al., 2013). Each environmental sound stimulus was cut to 950 ms in length. Furthermore, 16 disyllabic pseudowords were presented. The pseudowords were recorded by a native female speaker of German, were derived from existing German nouns, and were each cut to a length of 750 ms. The stimuli for this task are identical to previous experiments in auditory associative word learning (Cosper et al., 2020, 2022, 2024). The entire list of stimuli can be found in Supplementary Materials (A & B) and all files are additionally uploaded to the online repository. Each sound file was saved as a monaural .WAV file and digitized at 44100 Hz. During the experiment, all sound files were presented over two speakers on either side of the screen at a comfortable volume, which was indicated by each participant. The experiment, and thus the presentation of the stimuli, was conducted in a shielded EEG cabin, which had an additional benefit of blocking out background noise. The experiment was coded and presented with the Presentation software (Neurobehavioral Systems, Inc., Berkeley, CA, USA, version 24.007.19.23)

#### Episodic Short-Term Memory Recognition (STM) Tasks

For the auditory-nonverbal STM task, a total of 96 environmental sounds were presented. The sounds were acquired from databases of previous publications (Gygi & Shafiro, 2010; Norman-Haignere et al., 2015) or taken from freesound.org. The environmental sounds in this task were all distinctly different from the stimuli presented in the associative word learning task. A full list of all sounds and their sources can be found in the Supplementary Materials (C) and also be found on the online repository. Each sound was cut to exactly 1000 ms, saved as a monaural .WAV file, digitized at 44100 Hz, and presented over two speakers on either side of the screen at an individualized comfortable volume. The stimuli and the task were presented and conducted in a shielded EEG cabin using the same Presentation software.

For the auditory-verbal STM task, a total of 96 disyllabic German words were presented. Each word belonged either to the category of noun, adjective, adverb/preposition/pronoun, or verb with 24 words in each category. Word stems were only presented in a single form (i.e., in one of the four categories, but not in multiple categories). Each word was recorded by a female speaker of German, were cut to 1000 ms (including silence after the word), digitized at 44100 Hz, and saved as monaural .WAV files. A full list of the words and their English translations can be found in Supplementary Materials (D) and all files can be found on the online repository. The task was conducted in the EEG cabin and used the same Presentation software.

For the visual-nonverbal task, a total of 96 pictures were presented. All visual stimuli were taken from the Amsterdam Library of Object Images (Geusebroek et al., 2005). Each picture was presented in the center of the screen for 1000 ms. All visual stimuli were presented at a size of 768 x 576 pixels. A full list of visual stimuli can be found in Supplementary Materials (E) and in the online repository. All stimuli were presented on the screen in complete silence using the same Presentation software.

### 2.3 Procedure

The experimental procedure for each participant followed a strict structure over two visits to the lab. On the first visit, parents/guardians and participants were given details about the structure of both visits. Afterwards, both parents/guardians and participants read and signed consent forms, clarifying any questions that did not pertain to the goals of the study. Participants then completed the language development assessment test battery (SET 5-10; Petermann, 2012; Rißling et al., 2013) to conclude the first visit, which lasted for a total of 1 hour. During the second visit, participants were asked to complete the auditory associative word learning task and three short-term episodic recognition memory tasks. These were presented as playing four memory-style games on the computer situated in the EEG cabin. The tasks were presented in a fixed order to ensure that motivation was maintained until the end of the session (the order was determined through piloting). First, the AAWL task was completed. The second task was the auditory-nonverbal STM task, followed by the visual-nonverbal STM task. The fourth and final task was the auditory-verbal STM task. Between each task, the participants were offered a short break. This maintained motivation and active participation. The second visit lasted on average 90 minutes, at most 120 minutes. Each task is explained in detail below in accordance to the order of completion.

#### Language development assessment (SET 5-10)

In order to quantify the language abilities and development of each participant, a standardized diagnostic language development assessment was conducted with each participant – namely the SET 5-10 (Petermann, 2012). This assessment was created for German speaking children between the ages of 5;0 to 10;11 years old. The SET 5-10 test battery is a diagnostic tool used to gauge the language development of children and identify possible language processing issues. In its verification, the SET 5-10 can be conducted with children to diagnose language areas where processing and production may deviate from the standard due to, for example, language developmental delays, language impairments, learning disabilities, children with a migratory background, and children with brain injuries (e.g., aphasia). However, the use of this diagnostic language development assessment within the frame of this project was not to diagnose children or to identify any deviation from the typically developing cognitive span, but rather merely to quantify the language ability of each participant. This can be done as the subtests of the SET 5-10 correlate to assessments of language development and each subtest was validated in increments of 6 months between the age groups included in the test (cf. Petermann, 2012). As such, the use of the raw data from the SET 5-10 subtests yields itself as a measurement of language development. To further support this, the total scores of participants included in the final dataset of the two age groups in the current study were compared using an independent two-sample *t*-test and a corresponding Cohen’s d calculated in R (R Core Team, 2022) using the same method as the behavioral statistics described in section 2.5 below. Results indicated that 9-10-year-olds overall scored significantly higher than 5-6-year-olds, as shown by a two-sample independent *t*-test, *t*(-13.906) = 68.692, *p* < 0.001, with a large effect size (|*d|* = 2.90).

The results of this assessment were not provided to the participants or the families. This point was made clear to the parents and is printed explicitly in the signed consent forms. For the assessment, the participant sat at a table in a room with 1-2 experimenters (one to conduct the test, one to protocol the participant responses, some assistants conducted both tasks). The participants then completed 10 tasks with the experimenter. Participants were informed on how to conduct each task from the experimenter and given examples, as stated in the SET 5-10 guidelines (Petermann, 2012). Between each task, participants were offered a short break.

The SET 5-10 tested children’s language development and abilities in 7 areas, with a total of 10 specific assessments. The first assessment area was vocabulary with a picture-naming task. The second assessment area was semantic relations with a visual categorization task. Processing speed was the third assessment area with a timed visual search task. The fourth assessment area was language comprehension and was split into two tasks: an action sequence task and text comprehension task (the latter split into two assessment levels: 5 to 6-year-olds and 7 to 10-year-olds). Language production was the fifth area with two assessments: a picture story task and a sentence production task. The sixth area was for morphology and consisted of two assessments: a singular and plural production task and correct/incorrect sentence recognition task (the latter split into two levels: 5 to 6-year-olds and 7 to 10-year-olds). The seventh and final assessment area was auditory abilities with a pseudoword production task for 5 to 6-year-olds; however, the older children in the current study also completed this task. The experimenters completed a digital protocol for all answers. The language assessment took between 30-45 minutes in total.

#### AAWL Task

The sequential auditory associative word learning task was identical to previous iterations of the task (cf. Cosper et al., 2020, 2022, 2024), with minimal hardware and instructional differences. For the duration of the task, participants were seated in an EEG cabin at a desk with a computer screen, speakers on either side, and a keyboard in front of the participant. For this experiment, the participants also wore an EEG cap (see section 2.4 for detailed information). The participants were read the instructions for the experiment and were familiarized with the forced-choice button-press procedure for the testing phase. Participants were instructed to sit as still as possible and to pay attention to the sounds presented, as they would be asked questions about the sounds later. To ensure participation, an experimenter silently sat in the EEG cabin out of view of the participant. The experiment was divided into two phases: a training phase and a testing phase. The training phase consisted of 128 trials of sound-word combinations. Eight of the 16 sounds and words were allocated to the consistent condition, and the remainder 64 trials were allocated to the inconsistent condition. Consistent pairings were indicated by presenting the sound and the word consistently together over the course of the training phase 8 times for each combination, for a total of 64 trials. The inconsistent condition was comprised of 64 unique combinations of sounds and words, where each sound was paired with each pseudoword only once (for comparable training phase designs, see: Cosper et al., 2020, 2022, 2024; Friedrich & Friederici, 2008, 2011). The testing phase immediately followed the training phase, with a short, 30 second pause in between. The testing phase was comprised of 32 trials in two conditions, i.e., 16 matching pairings and 16 violated pairings. The matching pairings were the 8 sound-pseudoword combinations from the consistent pairings of the training phase, presented two times each, for a total of 16 trials. The violated pairings also used the 8 sounds and pseudowords from the consistent pairings, but were 8 novel pairings, which broke the expectation from the training phase. Each violated sound-pseudoword pairing was presented twice, for a total of 16 pairs. See Figure 1A for example for each of the conditions across the training phase and Figure 1B for an example of the testing phase in each condition.

**Figure 1:**
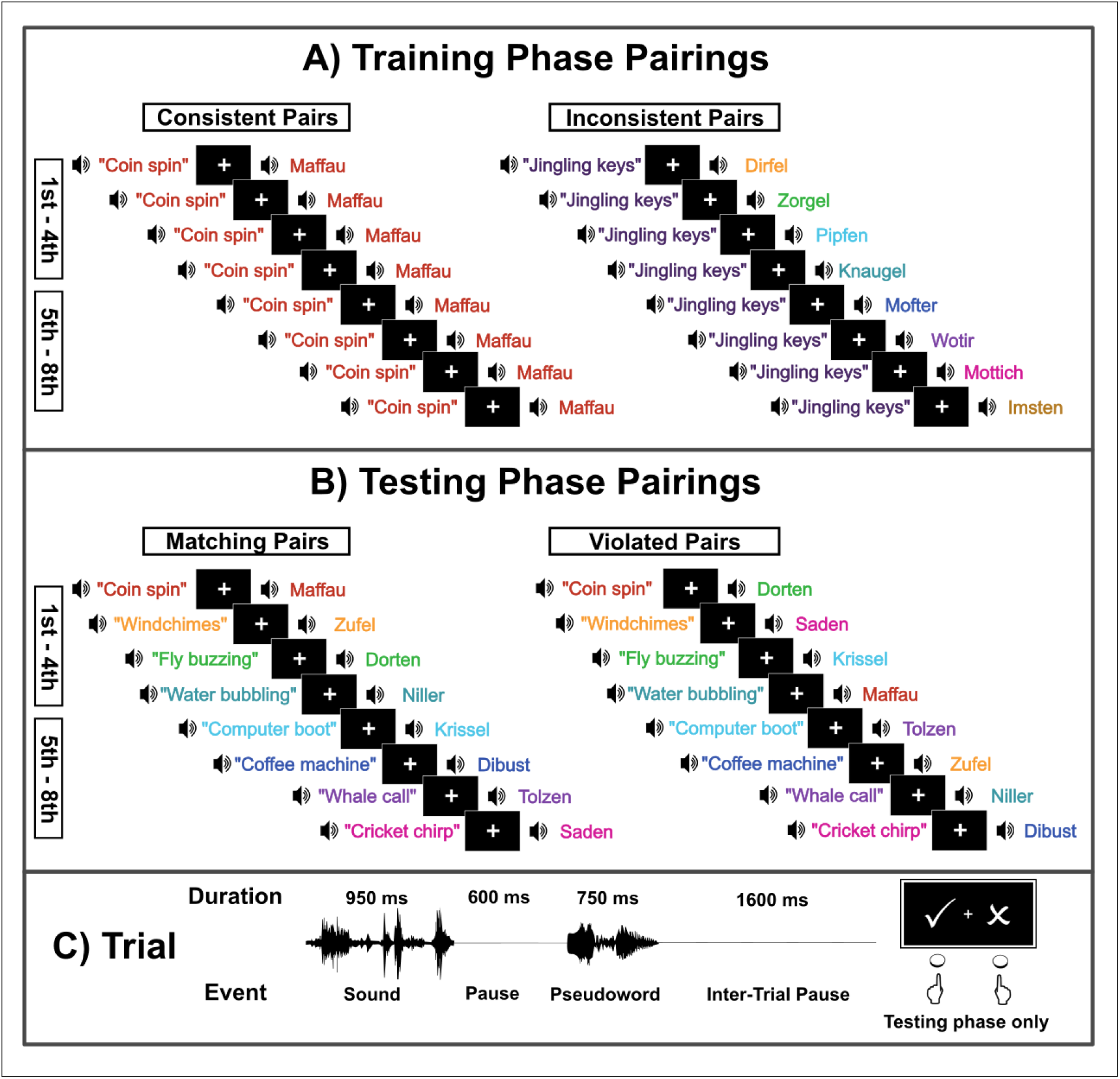
Experimental setup of the auditory associative word learning task. (A) Example of pairings of sound-pseudoword combinations for the training phase as presented to both age groups. (B) Example of pairings of sound-pseudoword combinations for the testing phase as presented to both age groups. (C) Trial structure for both phases, with the forced-choice button-press for the testing phase. Colors used for visual emphasis.

In order to account for possible sequence effects, eight separate lists of stimuli combinations and presentation orders were created. This counterbalanced the sound-pseudoword combinations and which sounds and pseudowords were allocated to consistent and inconsistent conditions in the training phase to ensure that underlying sound-symbolism effects were not facilitating any learning effects. Furthermore, presentation lists were inverted to reduce the likelihood of recency effects in learning. Participants were automatically allocated to specific randomized lists in accordance to their participant number. During the entirety of the experiment, a fixation cross was presented in the center of the screen.

The structure of each trial was identical for the training phase and for the testing phase. The environmental sound was presented at the beginning of each trial followed by an inter-stimulus pause of 600 ms (from offset of the sound to the onset of the pseudoword). After the inter-stimulus pause, the pseudoword was presented and was followed by an inter-trial pause of 1600 ms (see Figure 1C for the trial structure). To ensure that all stimuli were presented at the appropriate time, the Presentation Mixer was used (Neurobehavioral Systems, Inc., Berkeley, CA, USA, version 24.007.19.23). This resulted in all stimuli being presented within 1 ms of the intended time window and was measured by the uncertainty value in the log files. In cases of extended stimuli presentation latencies, the program automatically ended the experiment and the participant was removed from the final dataset – this only occurred twice in all participant measurements. During the testing phase, the participants were additionally asked to provide behavioral input in the form of forced-choice button-presses on the keyboard with accompanying pictorial indications on the screen (see Figure 1C for an example). After the inter-trial pause, participants were asked to judge if the presented sound-pseudoword combination was correct or incorrect. Assignment of left and right buttons for correct and incorrect judgements was counterbalanced in the randomization lists and was automatically applied according to participant number. In total, the experiment lasted 14.5 minutes. The Presentation files (including experiment file, PCL file, and scenario file) can be found on the online repository. After completion of this task, the EEG cap was removed. Participants were also allowed to wash their hair before continuing with the short-term episodic recognition memory tasks.

#### STM Tasks

Each of the three episodic STM recognition tasks (auditory-nonverbal, visual-nonverbal, and auditory-verbal) were presented in separate tasks. Visual-verbal STM was not included as reading is first expected to be learned in the first grade in Germany and not all participants had begun schooling at the time of testing. The structure of each STM task was identical (see Figure 2). Each task was presented to the participants as a game of memory. For each task, participants were presented stimuli in two phases: a presentation phase and a testing phase (cf. Cohen et al., 2009). In the presentation phase, 64 images, sounds, or disyllabic words were presented one time each. Each stimulus was presented for the same amount of time: 1000 ms. There was a pause between each stimulus of 1600 ms. Subsequently, participants entered the recognition phase. The recognition phase presented again 64 pictures, words, or sounds in the same manner and modality as the presentation phase, presented each only once. Half of the stimuli matched the presentation phase (old), while the other half was compiled of new stimuli. Each stimulus was presented for 1000 ms, as in the presentation phase, and was followed by an inter-trial pause of 1600 ms. After each stimulus presentation and inter-trial pause in the recognition phase, participants judged if the stimulus was the same as the presentation phase or different (old or new) via forced-choice button-press responses on the keyboard and accompanying pictorial indications on the screen (see Figure 2B for an example). Each test is presented in completion (presentation phase followed by testing phase) before advancing to the next test. These were conducted via the Presentation software while the participant is still in the EEG cabin. In order to avoid stimuli preference, presentation order preference, or other biases, the presentation of each phase was completely randomized within each condition, creating eight randomization lists in total. Left and right button-press responses of old and new were counterbalanced across the randomization lists. Allocation to a randomization list was automatic and determined by participant number. See Figure 2A for an example of the presentation phase for each STM task and Figure 2B for an example of the recognition phase of each STM task. For the auditory-nonverbal and auditory-verbal STM tasks, a fixation cross was always present on the screen. For the visual-nonverbal task, the fixation cross was presented during the inter-trial pauses. Each of the STM tasks took 11 minutes to complete.

**Figure 2:**
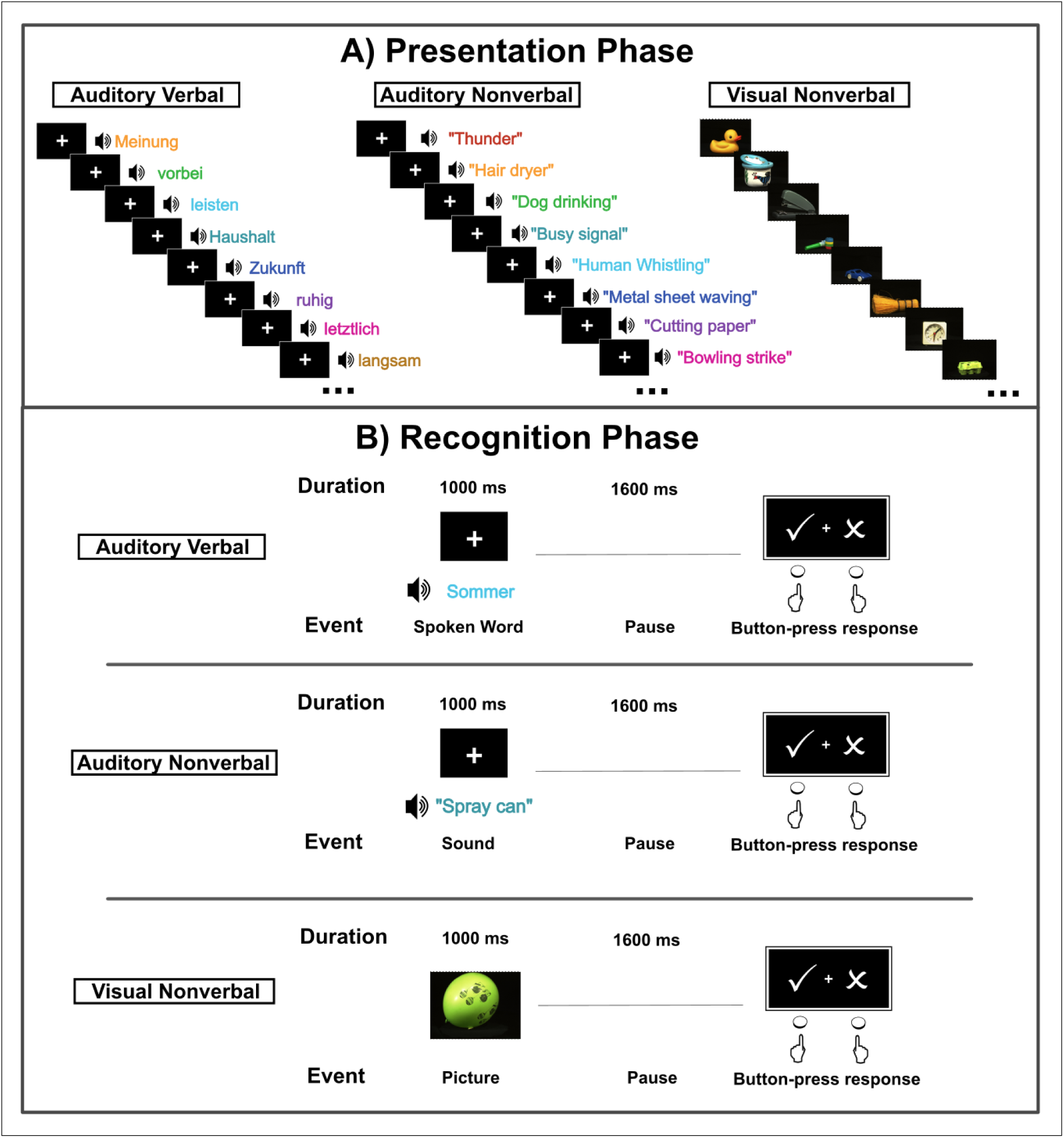
Experimental setup of the STM tasks. (A) Examples of presentations for the presentation phase of each modality. (B) Examples of the recognition phase for each modality. Each modality was presented in its own task. Colors used for visual emphasis.

### 2.4 EEG Processing

All data were collected at the Lifelab of the Technische Universität Dresden in a shielded room (EEG Cabin). EEG signals were measured using BrainAmp amplifier (Brain Products GmbH, Gliching, Germany) using Ag/AgCl electrodes at extended standard 10-20 positions in a custom Fast’n Easy BrainCap (sub-inion model; Brain Products GmbH, Gliching, Germany), with 60 equidistant sintered ring electrodes. For all electrode positions, see Supplementary Materials (F). Eye movement-related activities were measured by a singular monopolar electrooculogram (EOG) electrode under the left eye. All impedances were kept below 10 k Ω during the preparation process by means of a color-coded visualization on the screen, wherein contact between electrodes and the scalp was improved by adding gel or moving hair out of the way using a cotton swab (the process was given by Brain Products GmbH, Gliching, Germany). EEG data were continuously recorded using the BrainVision Recorder (Brain Products GmbH, Gliching, Germany, version 1.25.0202) with a sampling rate of 500 Hz, reference at Fpz, and ground located on the cap between electrodes AFz and Fz.

EEG pre-processing was conducted in MATLAB (The Mathworks Inc., Natick, MA, USA, version 2022b) using EEGLAB (Delorme & Makeig, 2004, version 14_1_1b) following the same protocol as Cosper and colleagues (2022, 2024). In a first step, a high-pass filter of 1 Hz (− 3dB, cutoff frequency of 1.38 Hz) and a low-pass filter of 30 Hz (− 3dB, cutoff frequency of 31.16 Hz) were applied to the continuous EEG data. Subsequently, the data was epochized with epochs time-locked to the onset of the pseudoword for a length of 1200 ms and a pre-stimulus baseline of -200 ms. After epochization, EEG data were re-referenced to averaged mastoids (scalp electrodes M1 and M2 – in this particular cap configuration, electrodes were labeled TP9 and TP10). Following, data were manually visually inspected for artifacts and any artifacts related to muscle movement, slow-drifts, or amplifier-blocking were removed. Afterwards, a semi-automatic independent component analysis (ICA) was applied to each dataset to correct eye-movement artifacts (reference and EOG electrodes were excluded from the ICA). A maximum of two components were removed (one for vertical eye-movements and one for horizontal eye-movements). After the ICA, the ICA weights of the 1 – 30 Hz filtered dataset were applied to a dataset with a high-pass filter of 0.3 Hz (− 3dB, cutoff frequency of 0.36 Hz) and a low-pass filter of 30 Hz (− 3dB, cutoff frequency of 31.16 Hz). Two separate filter settings were used for the ICA as artifact correction is best conducted with 1 Hz filtered data, but a 1 Hz high-pass filter is too strong for ERP statistical analyses (Klug & Gramann, 2021). This procedure is comparable to current ICA standards (Delorme, 2023). Subsequently, a separate manual visual artifact rejection was conducted with the 0.3 – 30 Hz dataset following the same guidelines as above. If necessary, electrodes were interpolated (maximum 6 channels per recording – 10% of channels). All statistical analyses were conducted using the 0.3 – 30 Hz filtered datasets.

In a last step, grand means were calculated across participants in each age group for each condition in the training phase and testing phase: consistent condition in the first half of the training phase (1^st^ – 4^th^ presentation of each pseudoword; 32 total), inconsistent condition in the first half of the training phase (1^st^ – 4^th^ presentation of each pseudoword; 32 total), consistent condition in the second half of the training phase (5^th^ – 8^th^ presentation of each pseudoword; 32 total), inconsistent condition in the second half of the training phase (5^th^ – 8^th^ presentation of each pseudoword; 32 total), matching condition of the testing phase (16 total) and the violated condition of the testing phase (16 total). As aforementioned, participants must have retained at least 50% of each trigger type after the ICA to be included in the final analysis. The average number of trials retained for each condition, the percentage of trials per condition retained, and their standard deviations for each age group can be seen in Table 1. All preprocessing scripts are available on the online repository.

### 2.5 Statistical Analysis

#### AAWL Task (EEG)

The statistical analyses of the EEG data were performed in accordance to previous studies in adult AAWL (Cosper et al., 2022, 2024). All statistical analyses for the EEG data of both age groups was conducted in MATLAB (The Mathworks Inc., Natick, MA, USA, version 2022b) using the FieldTrip toolbox (Oostenveld et al., 2011, version 22020104). For the analysis, a cluster-based permutation approach was taken by conducting dependent samples *t*-tests for each sampling point with the alpha threshold set at 0.05. In order for a significant sample to be included in the clustering algorithm, a minimum number of neighboring channels for the cluster-based permutation analyses was set to 2 channels. The triangulation method from FieldTrip was used to define neighboring channels. The Monte Carlo method (Maris & Oostenveld, 2007) of cluster-based permutation tests was applied with 1000 randomizations and an accompanying alpha threshold of 0.05. For the statistical analysis of the clusters, the dependent samples *t*-test (ft_statfun_depsamplesT) was used, a correction for multiple comparisons was applied over the cluster (cfg.correctm = ’cluster’), and cluster statistics were run with maxsum. Reference electrodes (M1 and M2) and the EOG electrode were excluded from the cluster-based permutation tests. For the analysis, two predetermined time windows were selected based on previous studies using the same task (Cosper et al., 2022, 2024): a typical N400 time window (400 – 800 ms post-stimulus) and a later time window (800 – 1200 ms post-stimulus). The later time window was selected, as younger children often have delayed N400-like effects (cf. Junge et al., 2021) and the inclusion of the later time window allowed for a more robust data-driven approach.

Statistical analyses of the EEG data were conducted in three separate comparisons per age group: comparing inconsistent – consistent waveforms in the first half of the training phase, comparing inconsistent – consistent waveforms in the second half of the training phase, and comparing violated – matching waveforms in the testing phase. All clusters within the N400 and the later time window are reported. Additionally, all trend-level differences within clusters (0.1< *p* < 0.5) are also reported and clearly labeled as trends in both the text and figure comments. Additional exploratory statistical analyses were conducted for each of the three analyses, comparing males and females (for both the 5-6-year-old group and the 9-10-year-old group) and musically trained children and non-musically trained children (for the 9-10-year-old group only). Sex differences were included to evaluate any possible differences in the neural processes during word learning, as sex was considered in the behavioral analyses and has been found behaviorally in both children and adults (Kaushanskaya et al., 2011, 2013). Musically trained 9-10-year-olds were also compared to non-musically trained 9-10-year-olds, as musical training has been shown to affect word learning processes in both children and young adults (Cosper et al., 2024; Dittinger et al., 2016, 2017, 2019). Results of these additional analyses and the accompanying ERP figures can be found in Supplementary Materials (G – I).

For all cluster-based permutation tests, reported time windows of clusters containing significant waveform differences are not determinations of absolute time windows of the effect, as cluster-based permutation tests are not suitable for effect latency determination (Sassenhagen & Draschkow, 2019). Reported results only described approximate time windows of effects and which electrodes belong to the cluster in which significant differences are found.

In order to compare EEG differences at the group levels (age differences, sex differences by age group, and musical training for the older children), we applied the lmeEEG pipeline (Visalli et al., 2024) in MATLAB (The Mathworks Inc., Natick, MA, USA, version 2022b). The lmeEEG pipeline is conducted for the first half of training, the second half of training, and the testing phase separately for each of the four comparisons (age, sex (younger children), sex (older children), and musical training in older children). This method first conducted a linear mixed model (LMM) for each channel and event-locked timepoint on trial-wise EEG data. For the LMMs, we used three separate formulas. For age, the formula of the LMM was: condition (inconsistent vs. consistent or violation vs. matching) x age (young vs. old) with subject as a random factor. For sex, each age group was tested separately with the following LMM formula: condition (inconsistent vs. consistent or violation vs. matching) x sex (male vs. female) with subject as a random factor. For musical training, the following formula was used only on the 9-10-year-old data: condition (inconsistent vs. consistent or violation vs. matching) x musical training (music vs. none) with subject as a random factor. The second step performed mass univariate linear regressions on marginal EEG data, where random-effect contributions were removed from the EEG data. In the final step, permutations were tested and applied threshold-free cluster enhancement; neighboring electrodes were defined by channel locations. Limitations on temporal onsets and offsets of significant differences also apply to this analysis due to the cluster-based permutation approach (cf. Sassenhagen & Draschkow, 2019). All MATLAB scripts (including instructions, script order, additional analyses scripts, FieldTrip settings, lmeEEG data preparation and pipeline analyses, and plotting scripts) are available on the online repository.

#### Behavioral Statistics: AAWL Task (behavioral), STM tasks, and Behavioral Correlations

The behavioral and STM data (auditory-nonverbal, visual-nonverbal, and auditory-verbal) were statistically analyzed for each age group and modality in R (R Core Team, 2022, version 4.2.1). In order to assess both the accuracy of responses in the AAWL tasks as well as in the STM tasks while also reporting on possible biases in forced-choice button-press responses, we chose a signal detection approach for the analysis of the data, unlike previous investigations where only response accuracy was presented (cf. Cosper et al., 2022, 2024). Button-press response data for each of the four tasks was automatically extracted from the Presentation logfiles of each participant and identified as a hit, miss, correct rejection, or false alarm based using R (R Core Team, 2022, version 4.2.1) and saved into age-group lists according to task. In a second step, d-prime analyses were conducted for each task of each age group individually. Here, we realized that many of the d-prime values could not be calculated due to strong biases in the response data. Any correction of the values, i.e., adding limitations to the calculation of d-prime where ceiling and floor performance was modified by ±0.5, would have greatly modified the data to an extent that might not reflect valid interpretations. As such, we decided not to normalize the hit-rates and the false-alarm rates and instead to calculate the corrected hit-rate for each participant individually, using the following formula (for similar uses of corrected hit-rate as a measure of recognition memory, see Schultz, Sommer, et al., 2022; Schultz, Yoo, et al., 2022):

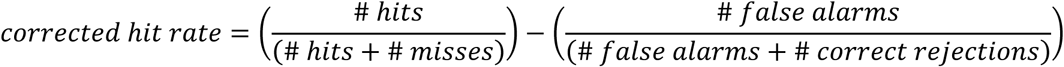

Though not normalized, the corrected hit-rate can be used to identify button-press biases (closer to 0 indicated a bias, closer to 1 indicated the absence of bias). In addition to bias, all statistical measures conducted on the corrected hit-rate were statistically proportional to the same statistical measures on response accuracy (with a chance level of µ = 0.5). However, for the sake of clarity, response accuracy ratio was calculated separately (# correct responses / total # questions) and is presented as a mean for each of the tasks per age group in percentages. Results for corrected hit-rate were assessed per task using a one-sample *t*-test against zero with an alpha threshold of 0.05. Cohen’s d was calculated for each one-sample *t*-test with the rstatix package (version 0.7.1) with µ = 0.0. Within each task, differences in corrected hit-rates between the two age groups were analyzed using an independent two-sample *t*-test and a corresponding Cohen’s d was calculated for each two-sample *t*-test.

Language development scores were calculated by the sum of the points accumulated by each participant in each of the 10 subtests of the SET 5-10. The values were not normalized in order to determine any diagnostic factors or risk factors, but were rather used as a quantitative measure of language development. SET 5-10 results were read into R and separated into the total value (SET score) and a value for each of the 7 language assessment areas: vocabulary, semantic relations (categories), visual processing speed, language comprehension, language production, morphology/syntax, and spoken pseudoword repetition. These scores were used for the correlation analysis and for the LMM as described below.

To examine the relationship between several variables, a Pearson correlation analysis was conducted. The correlation matrix included demographic variables (age at test, biological sex, music experience), language development (SET test scores), and corrected hit-rates across all conditions. False discovery rate (FDR) correction was applied to account for multiple comparisons. To display all significant correlations (α = .05; FDR corrected), the correlation matrix was visualized using the corrplot package (version 0.95).

To examine the effects of condition, age, sex, music experience and SET scores on the corrected hit-rate results, a linear mixed-effects model (LMM) was formed using the lme4 package (Bates et al., 2015, version 1.1.33). All variables were fixed effects while subject was included as a random intercept to account for repeated measures within participants. A null model including only the subject random intercept was specified to assess whether the chosen predictors significantly improve the model. In order to determine the if the fit model described more variance than the null model, AICs were compared and an ANOVA was run (see Supplementary Materials (J) for model comparisons). All R scripts for the statistical analyses, including all settings, are available on the online repository.

## 3 Results

### 3.1 Behavioral analyses

#### 5-6-year-olds

The behavioral word learning analysis revealed that 5-6-year-olds were only able to correctly identify matching and violated sound-pseudoword pairs in the testing phase at an average of 50.2 % of the time. The one-sample *t*-test indicated that the 5-6-year-olds were not above chance in identifying matching and violated pairings in the testing phase, *t*(45) = 0.16515, *p* = 0.867, with a negligible effect size (*d* = 0.024). The average corrected hit-rate for the AAWL task was 0.00408, indicating a strong bias in the button pressing. For the auditory-nonverbal STM, performance accuracy was 58.97% and the one-sample *t*-test revealed above chance performance in the identification of old vs. new environmental sounds *t*(45) = 6.7991, *p* < 0.0001, with a large effect size (*d* = 1.00). The average corrected hit-rate for the auditory-nonverbal task was 0.17935, indicating a bias in the button pressing. The auditory-verbal STM analysis yielded above chance performance in the identification of old vs. new spoken words, *t*(45) = 5.4868, *p* < 0.0001, with a large effect size (*d* = 0.809) with a performance accuracy of 57.58%. The average corrected hit-rate for the auditory-verbal task was 0.1515, indicating a bias in the button pressing. For the visual-nonverbal STM, performance accuracy was 73.54% and the one-sample *t*-test indicated above chance performance in identifying old vs. new pictures, *t*(45) = 8.4327, *p* < 0.0001, with a large effect size (*d* = 1.24). The average corrected hit-rate for the auditory-nonverbal task was 0.47079, indicating less evidence of bias in the button pressing. All results are summarized in Table 2, including confidence intervals.

**Table 2:**
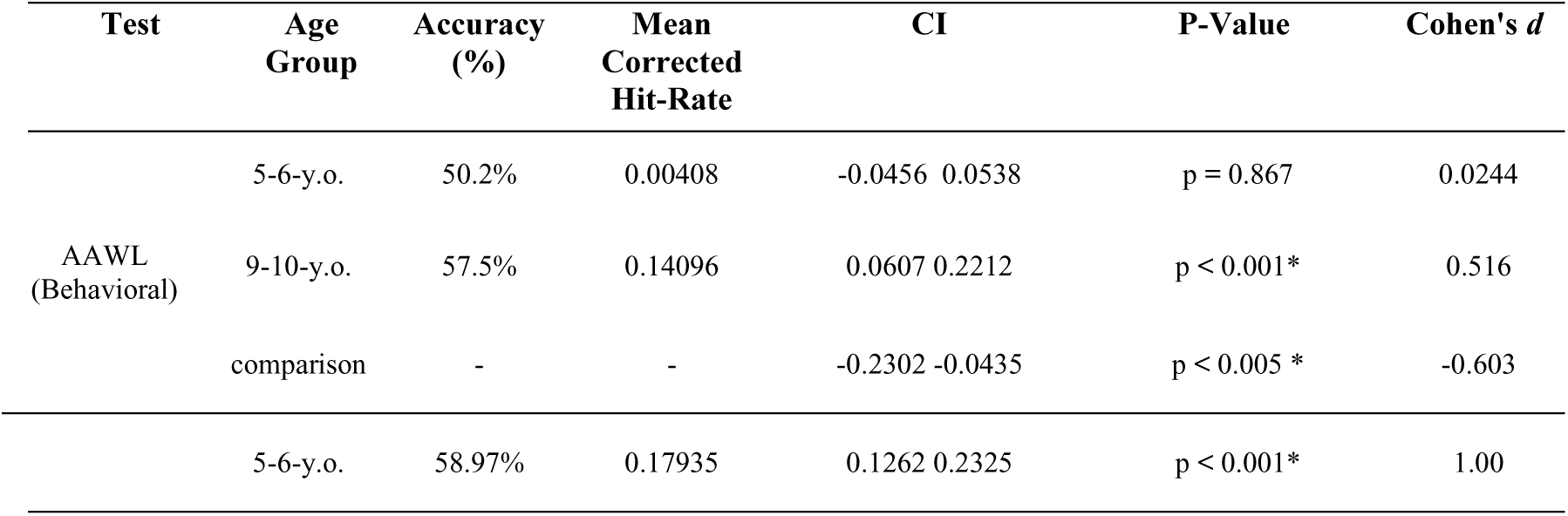
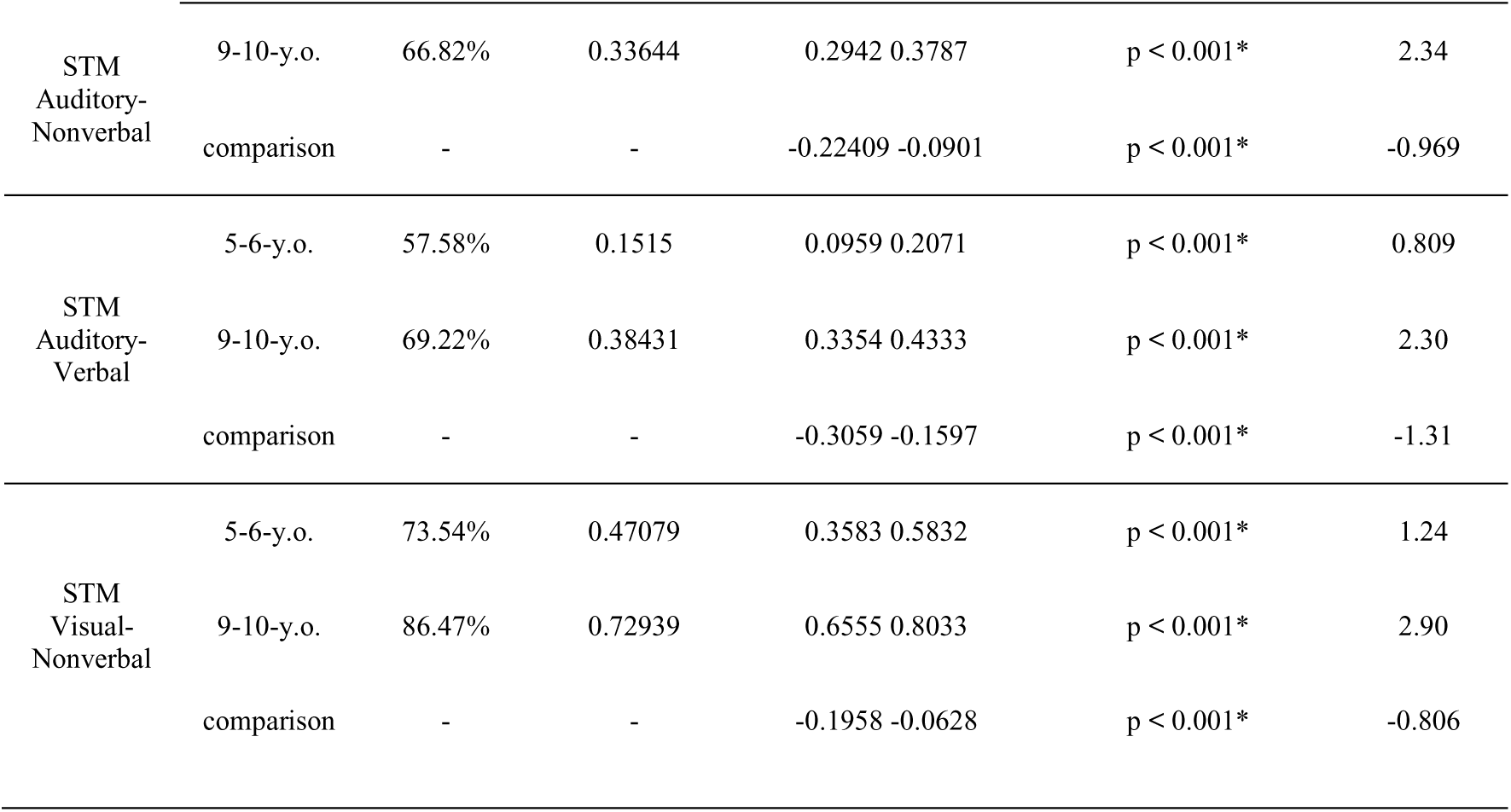
Behavioral results by test. Each test is further divided by age group and a comparison between the age groups. Confidence Intervals (CIs) are given for each one- and two-sample *t*-test. Significance is indicated by an asterisk (*). Mean corrected hit-rate is shown for each age group along with overall accuracy. Effect sizes are given for each *t*-test.

#### 9-10-year-olds

The behavioral word learning analysis revealed that 9-10-year-olds were able to correctly identify correct and incorrect sound-pseudoword pairs in the testing phase at an average of 57.5% of the time. The one-sample *t*-test indicated that the 9-10-year-olds performed above chance in identifying matching and violated pairings in the testing phase, *t*(46) = 3.5372, *p* < 0.001, with a moderate effect size (*d* = 0.516). The average corrected hit-rate for the AAWL task was 0.14096, indicating a bias in the button pressing. For the auditory-nonverbal STM, performance accuracy was 66.82% and the one-sample *t*-test indicated above chance performance in the identification of old vs. new environmental sounds *t*(46) = 16.036, *p* < 0.0001, with a large effect size (*d* = 2.34). The average corrected hit-rate for the auditory-nonverbal task was 0.33644, indicating a moderate bias in the button pressing. The auditory-verbal STM analysis yielded above chance performance in the identification of old vs. new spoken words, *t*(46) = 15.8, *p* < 0.0001, with a large effect size (*d* = 2.3) with a performance accuracy of 69.22%. The average corrected hit-rate for the auditory-verbal task was 0.38431, indicating a moderate bias in the button pressing. For the visual-nonverbal STM, performance accuracy was 73.54% and the one-sample *t*-test revealed above chance performance in identifying old vs. new pictures, *t*(46) = 19.872, *p* < 0.0001, with a large effect size (*d* = 2.9). The average corrected hit-rate for the visual-nonverbal task was 0.72939, indicating only a very slight bias in the button pressing. All results are summarized in Table 2, including confidence intervals.

#### Behavioral Results Comparison

In the AAWL behavioral task, 9-10-year-olds were able to identify matching and violated sound-pseudoword pairs significantly better than 5-6-year-olds, as shown by a two-sample independent *t*-test, *t*(76.545) = -2.9202, *p* < 0.005, with a moderate effect size (|*d|* = 0.603). In the auditory-nonverbal STM task, 9-10-year-olds outperformed 5-6-year-olds in their identification of old vs. new environmental sounds, as indicated by a two-sample independent *t-*test, *t*(86.197) = -4.6609, *p* < 0.0001, with a large effect size (|*d|* = 0.969). In the auditory-verbal STM task, 9-10-year-olds were significantly better than 5-6-year-olds in their identification of old vs. new spoken words, as shown by a two-sample independent *t-*test, *t*(89.322) = -6.327, *p* < 0.0001, with a large effect size (|*d|* = 1.3). For the visual-nonverbal STM task, 9-10-year-olds indicated a significantly better performance in identifying old vs. new pictures, as shown by a two-sample independent *t*-test, *t*(78.044) = -3.8705, *p* < 0.001, with a large effect size (|*d|* = 0.806). All results are summarized in Table 2, including confidence intervals.

### 3.2 EEG results (training phase)

#### 5-6-year-olds

For the first half of the training phase, in the whole sample no clusters were found which contained significant differences between inconsistent and consistent pairing for the 400 – 800 ms time window. For the 800 – 1200 ms time window, a negative cluster was found to have significant processing differences between inconsistent and consistent pairings from 822 – 1006 ms (*p* = 0.024; see Figure 3A). For exploratory sex differences, see Supplementary Materials (G).

**Figure 3:**
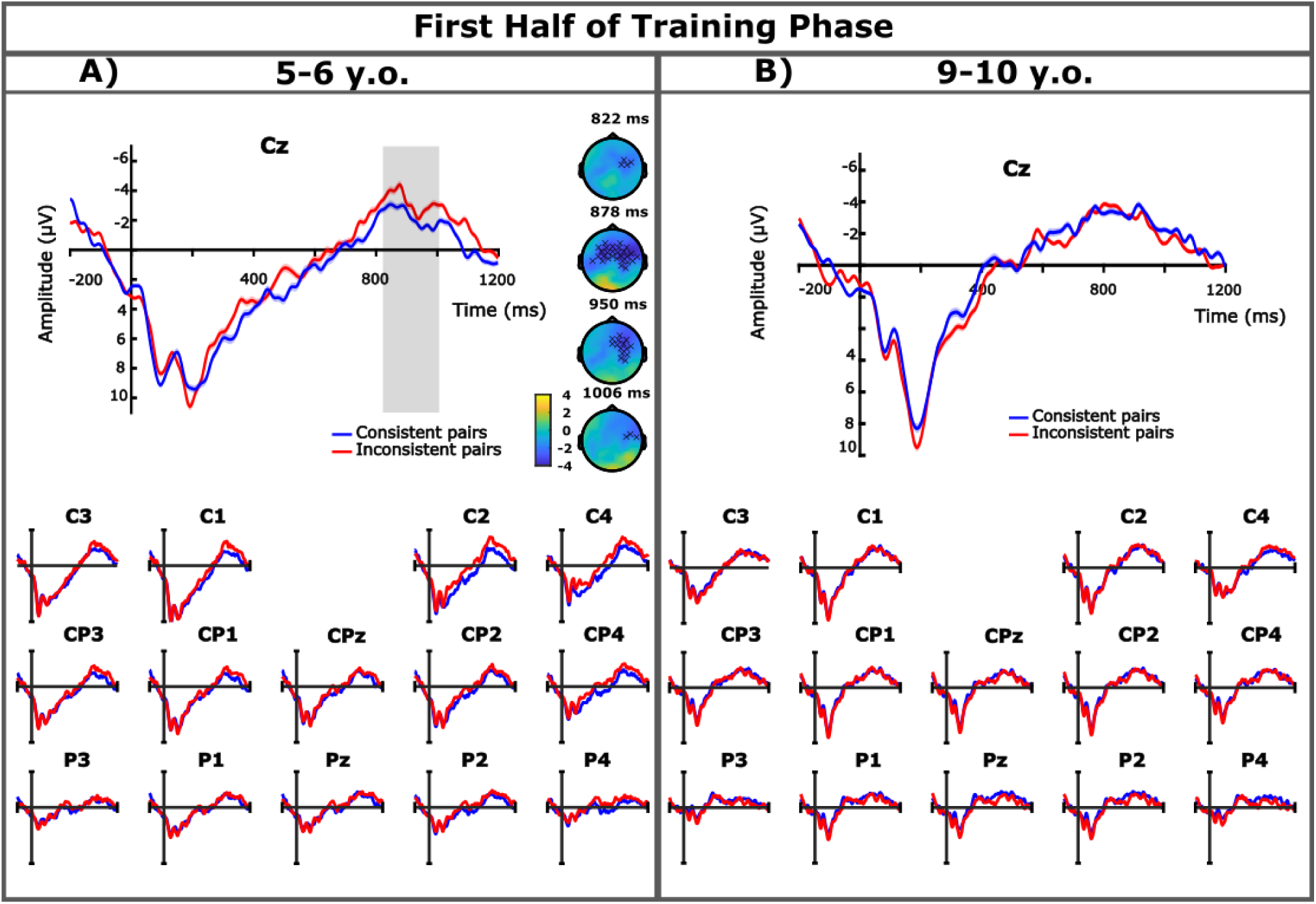
First half of the training phase results. ERPs are presented time-locked to the onset of the pseudoword for the first half of the training phase of each age group. The Cz electrode was selected as a representative electrode for both age groups, further centro-parietal electrodes are displayed below. The blue lines indicate the consistent pairs while the red lines indicate the inconsistent pairs. Gray bar indicates time window in which significant differences between the conditions were found. Topological difference maps from the cluster-based permutation test are presented only where significant differences were found and show the electrode distribution in the cluster over time (the X symbol indicates p < 0.05).

For the second half of the training phase, no clusters of significant processing differences were found in the 400 – 800 ms time window or in the 800 – 1200 ms time window. See Figure 4A for the ERPs. For exploratory sex differences, see Supplementary Materials (G).

**Figure 4:**
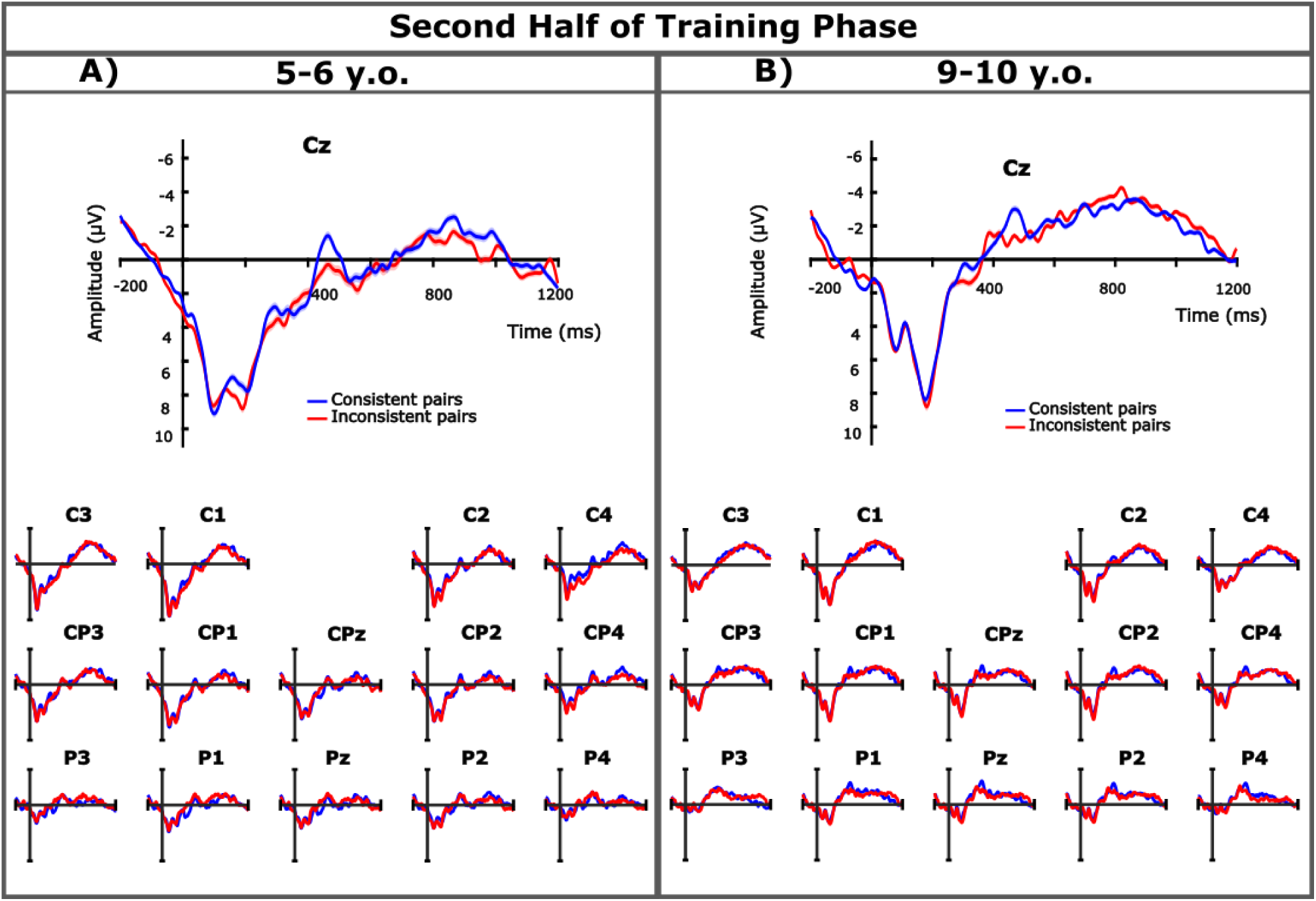
Second half of the training phase results. ERPs are presented time-locked to the onset of the pseudoword for the second half of the training phase of each age group. The Cz electrode was selected as a representative electrode for both age groups, further centro-parietal electrodes are displayed below. The blue lines indicate the consistent pairs while the red lines indicate the inconsistent pairs.

#### 9-10-year-olds

For the first half of the training phase, no clusters of significant processing differences between inconsistent and consistent pairings were found in either the N400 or the later time window (see Figure 3B). See Supplementary Materials (H) for exploratory sex differences and Supplementary Materials (I) for exploratory analysis on musical training.

For the second half of the training phase, no clusters with significant processing differences between inconsistent and consistent pairings were found for the 400 – 800 ms time window or for the 800 – 1200 ms time window (see Figure 4B). For exploratory analysis of sex differences, see Supplementary Materials (H). For exploratory analysis of musical training, see Supplementary Materials (I). All group differences are presented in Section 3.4 below.

### 3.3 EEG results (testing phase)

#### 5-6-year-olds

For the testing phase, no clusters of significance differences between violated and matching pairings were found for the 400 – 800 ms time window or for the 800 – 1200 ms time window (see Figure 5A). For the exploratory analysis of sex differences, see Supplementary Materials (G).

**Figure 5:**
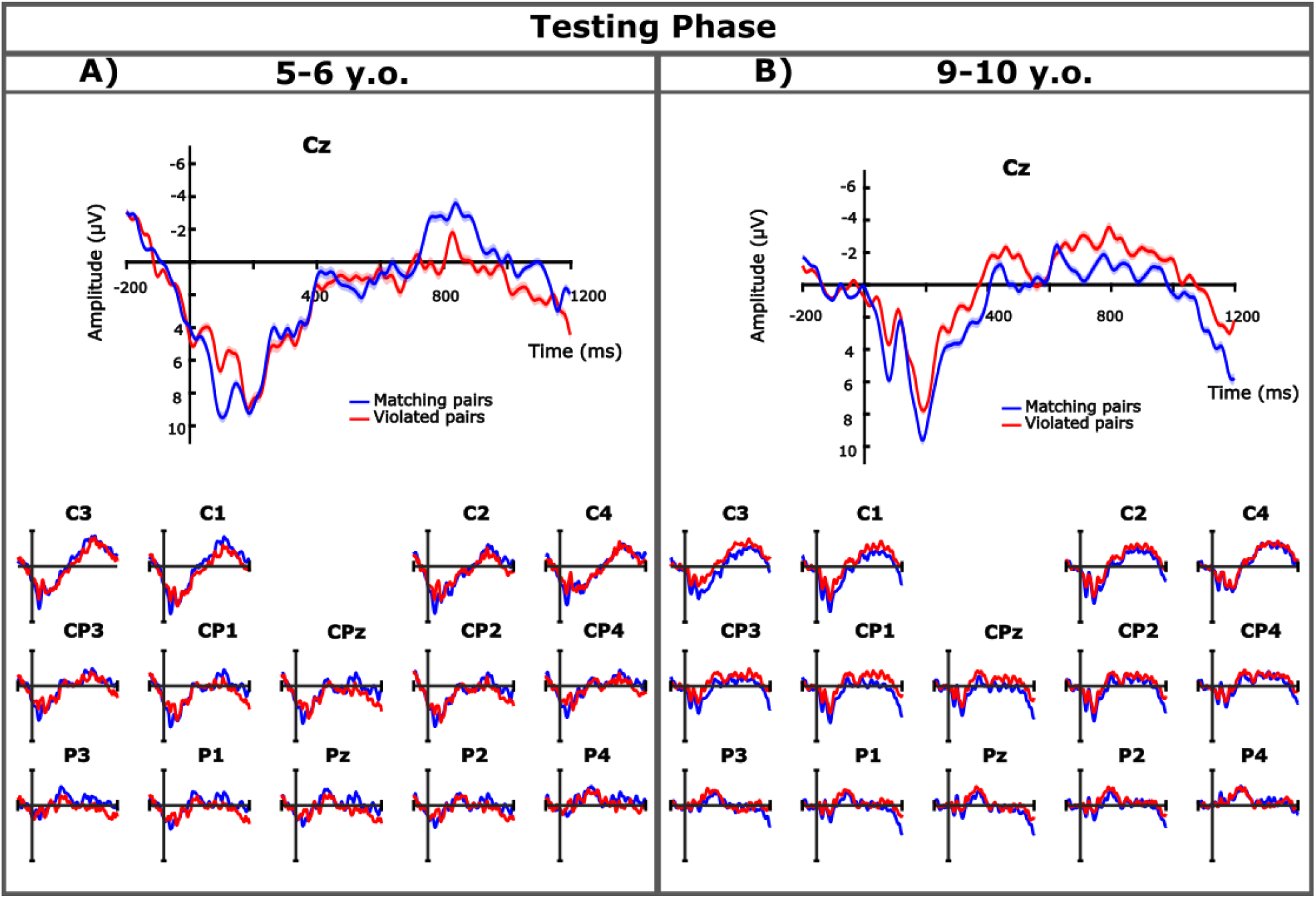
Testing phase results. ERPs are presented time-locked to the onset of the pseudoword for the testing phase of each age group. The Cz electrode was selected as a representative electrode for both age groups, further centro-parietal electrodes are displayed below. The blue lines indicate the matching pairs while the red lines indicate the violated pairs.

#### 9-10-year-olds

For the testing phase, no clusters with significant processing differences between violated and matching pairings were found for the N400 time window or the later time window (see Figure 5B). Furthermore, For the exploratory analysis of sex differences, see Supplementary Materials (H). For the exploratory analysis of musical training see Supplementary Materials (I). All group differences are presented in Section 3.4 below.

### 3.4 EEG group-level comparisons

Group level comparisons were conducted using the lmeEEG pipeline (Visalli et al., 2024). To this end, we presented all main effects and interactions in Table 3 below for the first half of the training phase, the second half of the training phase, and the testing phase. The main effects of condition and group comparison (age, sex, or musical training) were not relevant for this analysis, as condition was tested at the age level in sections 3.2 and 3.3 above for the exploratory analyses sex and musical training within the age groups presented in Supplementary Materials (G and H) and Supplementary Materials (I) in the cluster-based permutation tests. Age, sex, and musical training as main effects were not relevant, as they did not consider the condition of the EEG data (consistent vs. inconsistent or matching vs. violation). The interaction between condition and either age, sex, or musical training were of interest as all relevant parameters have been considered. All significant time windows are presented, even those outside of the N400 (400 – 800 ms) and later (800 – 1000 ms) time windows considered in sections 3.2 and 3.3 above and in the Supplementary Materials (G-I). Directionality of significant interaction differences were also noted in Table 3. Electrode clusters included in significant interactions (condition x group comparison) can be found in Figure 6.

**Table 3:** Group comparisons for EEG data. Each group is defined by age, sex (within a single age group) and musical training (for older children). Training 1 refers to the first half of the training phase while Training 2 refers to the second half of the training phase. Condition refers to inconsistent pairings vs. consistent pairings (in the training phase) or violated vs. matching parings in the testing phase. Group comparison is either age (older vs. younger), sex (male vs. female), or musical training (music vs. none). Interactions are between condition and the group comparison factor. All comparisons include subject as a random factor. Clusters containing significant differences in the EEG are indicated by a time window given in which the differences are found (in all cases *p* < 0.05, relevant results bolded for emphasis; *ns* = no clusters containing significant differences). Interpretations are given only for the interactions to indicate directionality.

| Group-Level | Experiment Phase | Condition | Group Comparison (Age, Sex, or Musical Training) | Interaction (Condition x Group Comparison) | Interpretation of Interaction |
| --- | --- | --- | --- | --- | --- |
| **Age (Older vs. Younger)** | Training 1 | ns | ns | ns | - |
|  | Training 2 | ns | ns | ns | - |
|  | Testing | ns | ns | ns | - |
| **Sex (Younger Male vs. Younger Female)** | Training 1 | 602 – 666 ms<br>766 – 856 ms | ns | ns | - |
|  | Training 2 | ns | 492 – 542 ms | ns | - |
|  | Testing | ns | ns | ns | - |
| **Sex (Older Male vs. Older Female)** | Training 1 | 162 – 266 ms | ns | **152 – 338 ms** | Early relative negativity larger in male participants than in female participants (outside of N400 time windows) |
|  | Training 2 | ns | ns | ns | - |
|  | Testing | ns | ns | ns | - |
| **Musical Training (Music vs. None)** | Training 1 | ns | ns | ns | - |
|  | Training 2 | 406 – 508 ms | 316 – 456 ms | **326 – 554 ms** | Relative positivity (LPC) larger in musical training group than in group without musical training |
|  | Testing | ns | 304 – 504 ms<br>524 – 608 ms<br>614 – 748 ms | ns | - |

**Figure 6:**
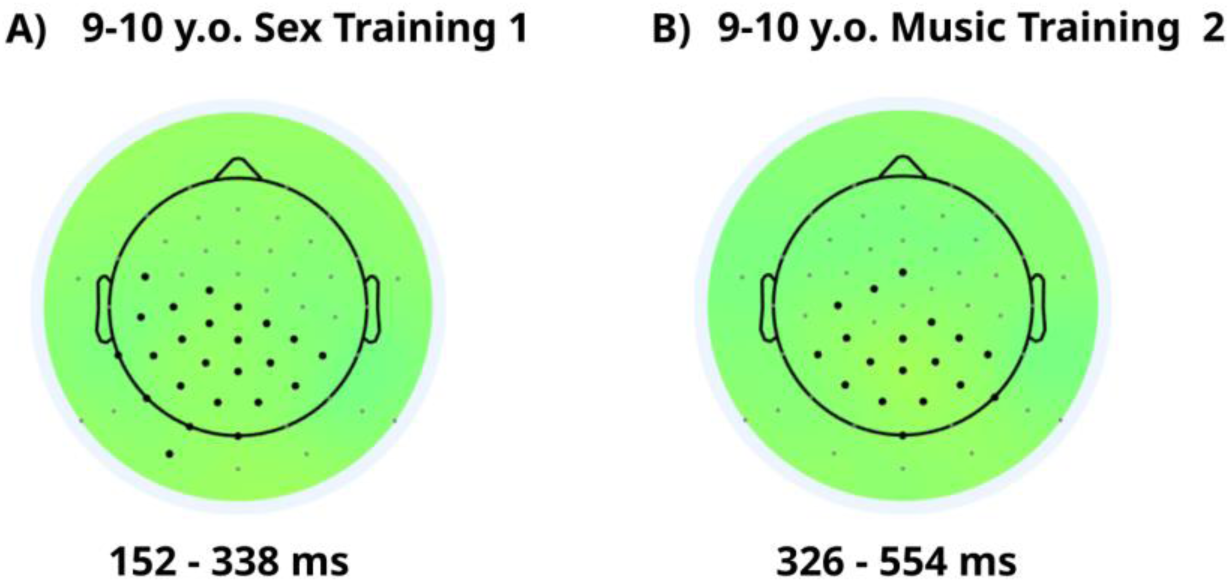
Electrode distribution in clusters of significant condition x group comparison differences. Training 1 refers to the first half of the training phase while Training 2 refers to the second half of the training phase. A. shows the cluster for the older children condition x sex differences found in the 152 – 338 ms time window. B. shows the cluster for the older children condition x musical training found in the 326 – 552 ms time window. Color of the head forms is a standard topoplot zero color. Electrode positions with labels can be found in Supplementary Materials (F).

### 3.5 Behavioral correlations with episodic memory measures

In order to determine if the LMM with fixed-factors explained more variance in the data than a model excluding fixed-factors, a two-way ANOVA was run, resulting in higher efficacy for the model with fixed-factors (χ^2^ = 267.02, *Df* = 8, *p* < 0.001). For more information on the model comparisons, see Supplementary Materials (J). The fixed-factors LMM yielded significant results between the STM tasks against the AAWL behavioral task. Specifically, the auditory-nonverbal STM task (*β* = -0.332, *SE* = 0.03, *t* = -11.266, *p <* 0.001), the auditory-verbal STM task (*β* = -0.528, *SE* = 0.03, *t* = -17.906, *p <* 0.001), and the visual-nonverbal STM task (*β* = -0.343, *SE* = 0.03, *t* = -11.619, *p <* 0.001) significantly affected the overall increase in behavioral performance, but the AAWL performance did not. Of the additional factors, only language development (SETscore; *β* = 0.004, *SE* = 0.001, *t* = 3.333, *p* = 0.001) was a significant factor for increased behavioral performance, while the other factors were not, including group (*β* = 0.093, *SE* = 0.115, *t* = 0.810, *p* = 0.418), age (*β* = -0.022, *SE* = 0.035, *t* = -0624, *p* = 0.533), sex (*β* = 0.051, *SE* = 0.031, *t* = 1.648, *p* = 0.1), and musical training (*β* = 0.021, *SE* = 0.038, *t* = 0.563, *p* = 0.574). See Table 4 to see all fixed-factors and random effects.

**Table 4:**
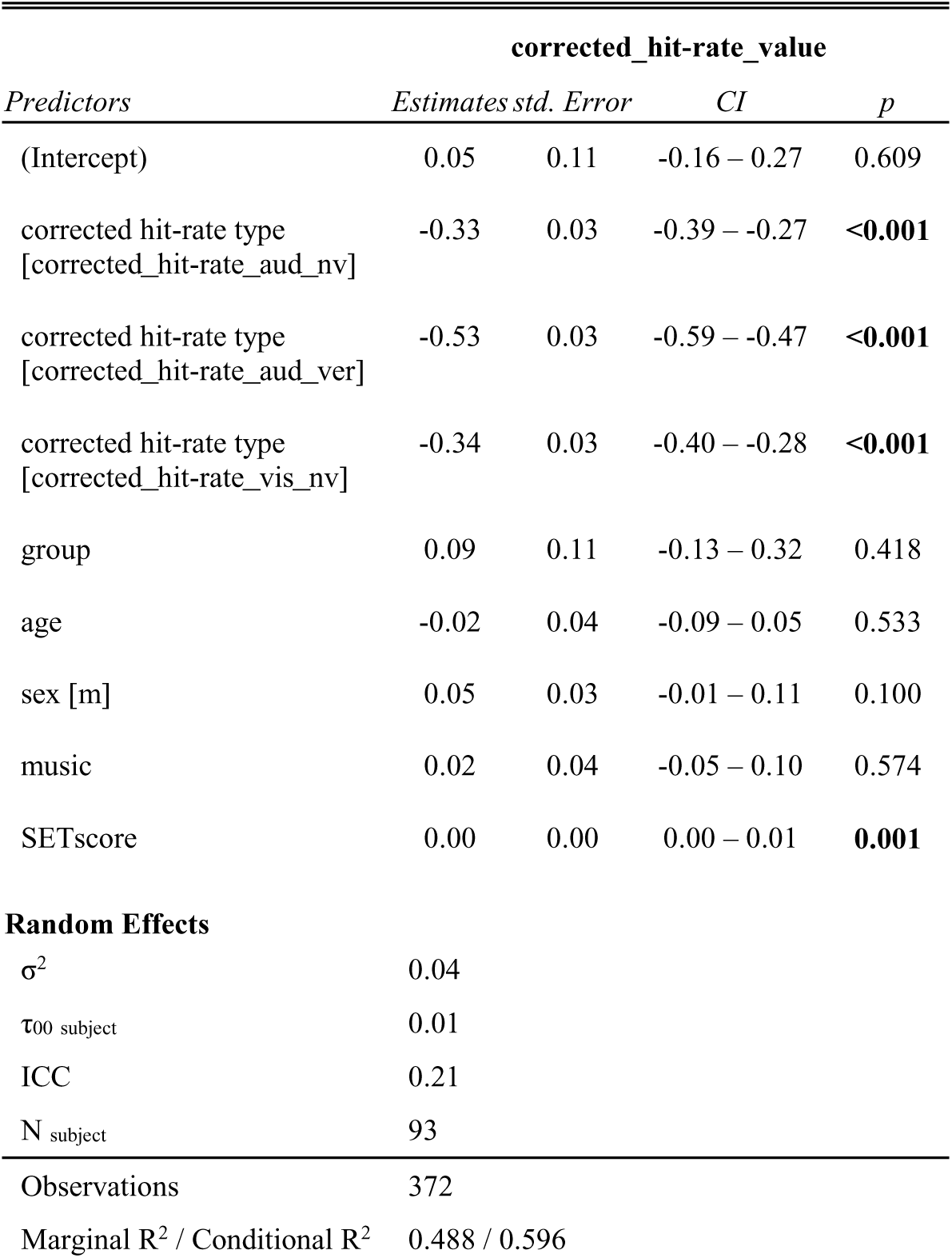
LMM results of the behavioral data for each of the predictors. Brackets indicate which predictor variable was compared to the reference variable (corrected_hit-rate_aawl and female). Random effects are listed below model. Significant values are bolded.

| *Predictors* | **corrected_hit-rate_value** |  |  |  |
| --- | --- | --- | --- | --- |
|  | *Estimates* | *std. Error* | *CI* | *p* |
| (Intercept) | 0.05 | 0.11 | -0.16 – 0.27 | 0.609 |
| corrected hit-rate type<br>[corrected_hit-rate_aud_nv] | -0.33 | 0.03 | -0.39 – -0.27 | **<0.001** |
| corrected hit-rate type<br>[corrected_hit-rate_aud_ver] | -0.53 | 0.03 | -0.59 – -0.47 | **<0.001** |
| corrected hit-rate type<br>[corrected_hit-rate_vis_nv] | -0.34 | 0.03 | -0.40 – -0.28 | **<0.001** |
| group | 0.09 | 0.11 | -0.13 – 0.32 | 0.418 |
| age | -0.02 | 0.04 | -0.09 – 0.05 | 0.533 |
| sex [m] | 0.05 | 0.03 | -0.01 – 0.11 | 0.100 |
| music | 0.02 | 0.04 | -0.05 – 0.10 | 0.574 |
| SETscore | 0.00 | 0.00 | 0.00 – 0.01 | **0.001** |
| **Random Effects** |  |  |  |  |
| $\sigma^2$ | 0.04 | | | |
| $\tau_{00}$ subject | 0.01 | | | |
| ICC | 0.21 |  |  |  |
| N<sub>subject</sub> | 93 |  |  |  |
| Observations | 372 |  |  |  |
| Marginal R<sup>2</sup> / Conditional R<sup>2</sup> | 0.488 / 0.596 |  |  |  |

Figure 7 shows the Pearson correlation matrix (FDR corrected) among all behavioral tasks, demographic information, musical training, and language development. From the correlation matrix, auditory-verbal STM performance was negatively correlated to musical training (*r*(91) = -.58, *p* <0.05), indicating higher recognition memory for non-musically trained participants. Furthermore, higher performance on the visual-nonverbal STM task was positively correlated with behavioral performance on the AAWL task (*r*(91) = .82, *p* <0.01). Language development was also positively correlated with group (*r*(91) = .91, *p* <0.01) and age at test (*r*(91) = .94, *p* <0.01), while negatively correlated to biological sex (*r*(91) = -.70, *p* <0.01), indicating that language development indexes were higher for female participants. For a complete table of all correlations and their *p*-values (FDR corrected), see Supplementary Materials (K).

**Figure 7:**
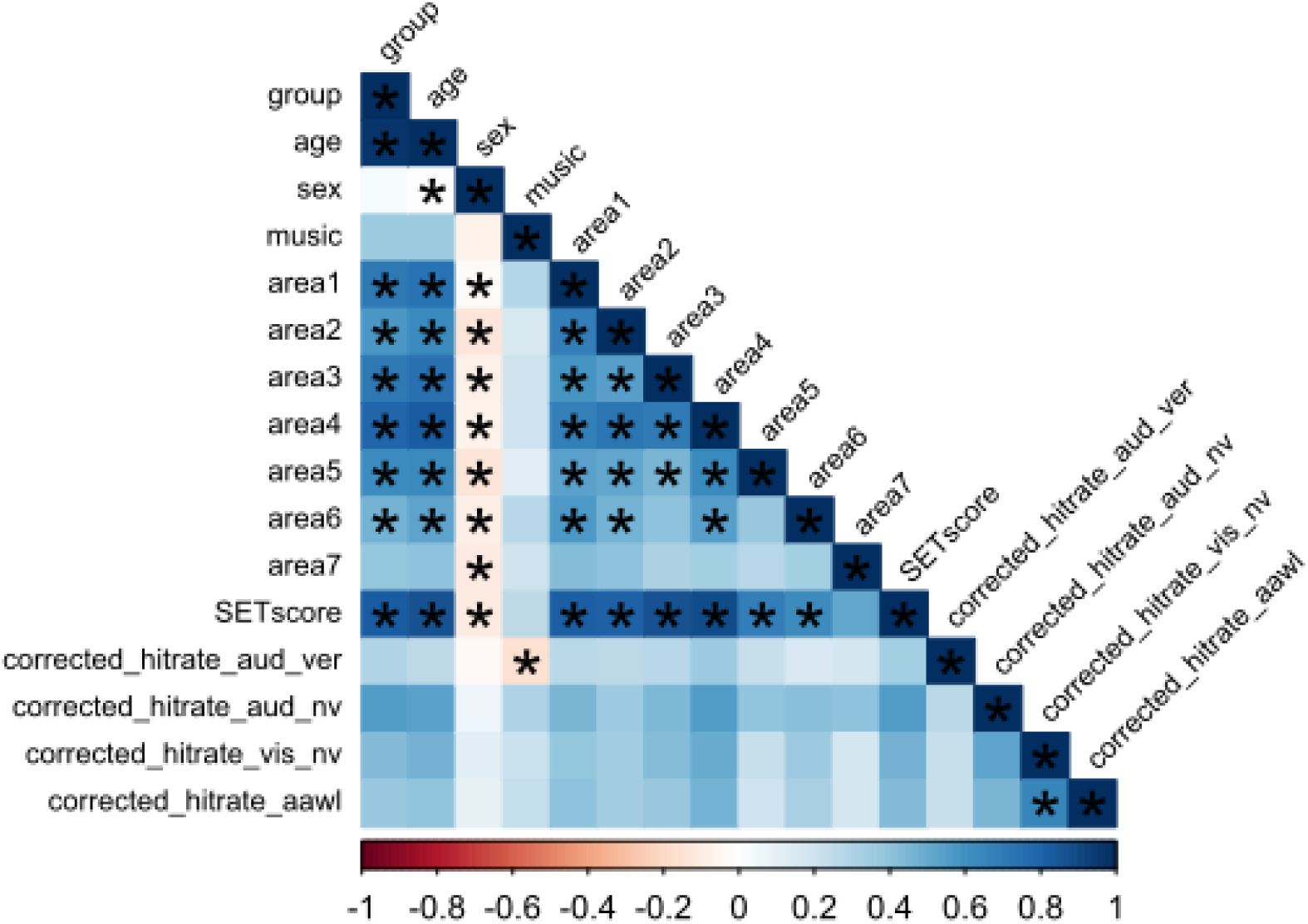
Pearson correlation matrix of behavioral tasks, demographic information, musical training, and language development. Group: 1(5-6-years-old), 2(9-10-years-old); age: actual age at time of test; sex: 1(female), 2(male); music: 0(no musical training), 1(musical training); area 1: Vocabulary; area 2: Semantic relation; area 3: (Visual) Processing speed; area 4: Language Comprehension; area 5: Language Production; area 6: Morphology/Syntax; area 7: Auditory Abilities (spoken pseudoword repetition); SETscore: total score among all 7 areas. For the corrected hit-rate values, aud-ver: auditory-verbal STM, aud_nv: auditory-nonverbal STM, vis_nv: visual-nonverbal STM; aawl: AAWL behavioral task. An asterisk (*) denotes significant correlation (α= 0.05). All correlations include FDR corrections. The color denotes the direction of correlation: red denotes a negative correlation and blue denotes a positive correlation.

## 4 Discussion

The aim of the present study was to investigate auditory associative word learning in 5-6-year-olds and 9-10-year-olds and to explore effects of short-term memory and language development on associative word learning with environmental sounds and spoken pseudowords. Our first hypothesis was that both age groups would be able to successfully map novel pseudowords onto environmental sounds both behaviorally and evidenced by the elicitation of the N400. We were unable to confirm this hypothesis in its entirety – only the older children indicated behavioral evidence of word learning (details below in section 4.1). Our second hypothesis stated that the N400 would present itself differently in younger and older children in the testing phase, with younger participants eliciting larger and earlier N400 effects, more similar to infant-like learning (Cosper et al., 2020) and that behavioral effects would be larger for younger children. This hypothesis was not confirmed. For our exploratory analyses, we assessed short-term memory and language development in both groups. We found that with age, language development and all forms of short-term episodic recognition memory improved substantially. We additionally explored the interplay between auditory associative word learning, language development, and short-term memory. Here, we found a positive correlation between visual-nonverbal STM and auditory associative word learning. All of our findings and additional explorative analyses are presented and discussed in detail below. We first addressed auditory associative word learning (training and testing phase results), including exploratory sex differences and exploratory musical training differences within each age group, followed by the exploratory investigation of STM and language development results. Finally, we discussed the limitations of the current set of experiments and outlook.

### 4.1 Auditory Associative Word Learning

#### 4.1.1 Training Phase

While we made no specific hypotheses about the training phase, it is important to address the results. Overall, 5-6-year-olds indicated a late negativity in the first half of the training phase, indicating an early sensitivity to consistent sound-pseudoword pairs, although no age-level differences were found in the EEG data for the training phase. When comparing sex differences, processing differences between consistently paired and inconsistently paired sounds and pseudowords in the form of an N400-like effect can be seen in the early time window for males, and in the early and late time windows for female participants, although no differences between 5-6-year old males and 5-6-year-old females were found at the group-level. Similar results for the first half of the training phase have been found in 10-12-month-old infants (Cosper et al., 2020). Cosper and colleagues (2020) found a pairing consistency effect in the first half of the training phase between inconsistently and consistently paired sounds and pseudowords. Their interpretation of the effect, despite opposite polarity and the heavy focus on the second half of the training phase, indicated the pairing consistency effect to be an indication of successful mapping of consistently paired items. Despite processing differences in the first half of the training phase in the form of a late N400-like effect (for N400-like latency justifications, see Junge et al., 2021), the second half of the training phase did not indicate any processing differences at the group level or in the analyses of male or female participants, indicating early processing differences or biases that did not lead to concrete learning effects during the course of the training phase.

For the 9-10-year-old group, there were no indications of processing differences in the first half or the second half of the training phase for the group level, for the analyses of males and females, or for non-musically trained children. However, differences were found early in the first half of the training phase between male and female older children at the group level (152-338 ms), suggesting early processing differences or biases between male and female older children. However, these differences did not persist in the second half of the training phase for the older children. Furthermore, significant differences were found in the second half of the training phase between the musically-trained children and children without musical training (between 326-554 ms). The musically trained older children revealed significant processing differences in the second half of the training phase; however, with reversed polarity – a relatively more positive waveform for inconsistent pairings than for consistent pairings. Similar effects have been found in the testing phase with auditory associative word learning in adult musicians (Cosper et al., 2024), which have been argued to mirror the positive component (LPC) effects of musicians distinguishing between congruent and incongruent musical melodies and rhythms (e.g., Besson & Faïta, 1995; Calma-Roddin & Drury, 2020; Featherstone et al., 2013; Miranda & Ullman, 2007; Steinbeis et al., 2006).

In the context of auditory associative word learning, infants have shown processing differences in both the first and second half of training (Cosper et al., 2020) while adults have shown some instances of processing differences in the second half of training with sequential stimulus presentation (Cosper et al., 2022), but not consistently under various circumstances including musician participants, athlete participants, or when stimuli are presented in reverse presentation order (Cosper et al., 2024). As such, the late N400-like effects in the first half of the training phase for the 5-6-year-olds reflected more the infant-like processing effects while the null-results of the 9-10-year-old group, including the LPC-like effect for the musically trained group in the second half of the training phase, paralleled the adult-like difficulties in processing of adult populations (Cosper et al., 2022, 2024). The presence of the LPC in the second half of the training phase suggests that even early musical training affects how the brain implicitly pairs auditory objects and spoken pseudowords.

Alternative interpretations for the N400-like effect for the younger children include heightened initial processing differences between consistent and inconsistent pairs, which may have tapered off due to memory-related factors of memory capacity, memory exceeding available cognitive resources, or cognitive load in the second half of the training phase, inhibiting overall learning (cf. Baddeley, 2012; Hubbard, 2010). Furthermore, the results of the musically trained 9-10-year-olds may not evidence lexico-semantic learning in the training phase. As indicated by Cosper and colleagues (2024), the LPC-like effect does not necessarily imply semantic processing. Instead, the LPC-like effect was interpreted in adults to be a form of implicit processing of congruent and incongruent auditory information unique to musicians (Besson & Faïta, 1995; Calma-Roddin & Drury, 2020; Featherstone et al., 2013; Miranda & Ullman, 2007). However, Kutas and Hillyard (1980) have further described the inversed-polarity N400-like effect in word learning situations not as a unique processing feature of musical training, but rather merely as a processing of physical differences between auditory stimuli of violated pairs and consistent pairs and not lexico-semantic differences between the two conditions. To what extent this inverted, relatively positive-going effect can be considered as a word learning effect is uncertain, as Cosper and colleagues (2020) considered the pairing consistency effect – a more positive-going wave form for inconsistent than consistent pairings during training – for infants to be a plausible indication sound-pseudoword association over time. Further testing is needed in order to determine how to interpret the more positive differences in waveforms in infant, child, and adult associative word learning.

#### 4.1.2 Testing Phase

For the testing phase, the 5-6-year-old children did not indicate above-chance behavioral word learning effects via button-pressing; however, the 9-10-year-old children showed significant above-chance learning effects in the behavioral measure. Further analysis confirmed that the older age group significantly outperformed the younger age group. There was no significant correlation between musical training, sex, or age and auditory associative word learning outcome as indicated by the button-press reaction. This did not confirm our hypothesis that younger children would evidence higher levels of learning than older children. These results are also subject to interpretation. Given the high button-press bias for the corrected hit-rate of the 5-6-year-olds in the AAWL task, it is possible that the associative word learning task may have been especially difficult for the younger children. This bias is seen in the GLM comparison, where all three episodic STM tasks significantly contributed to higher corrected hit-rates, while the AAWL task was not a significant factor. Furthermore, the experimental setup with EEG may have also been a contributing factor. The children only wore the EEG cap for the AAWL task, thus were asked to sit as still as possible; while the cap and all electrodes were removed for the STM tasks. Moreover, the order of the tasks may have also had an effect on learning outcome. As the tasks were performed in a fixed order with the AAWL task as the first task, the participants, particularly the younger participants, may have had initial difficulty with the button-pressing, which improved over time.

Unlike the ERP analysis of the training phase, neither the testing phase for the younger nor older children yielded any N400-like effects. Furthermore, there were no significant differences between the age groups. The further analysis between male and female participants for both age groups yielded similar non-significant results. However, the non-musically trained 9-10-year-olds indicated a trend-level N400 effect in the early time window. These differences, however, were not significant in comparison to the musically-trained older children at the group-level. While this can be a suggestion of possible associative word learning in the older, non-musically trained group, this trend-level effect was not seen in any of the further analyses. For adult populations, significant behavioral results were found when pseudowords were presented before sound stimuli with only trend-level N400-like effects in the testing phase (label-object-ordering group; Cosper et al., 2024), it has been hypothesized that this may have been due to the small sample size of the adult group. When comparing the behavioral effect sizes of the adult group (label-object-ordering) to the 9-10-year-old-group, the adults have a large effect size while the 9-10-year-olds only indicated a moderate effect size (cf. Cosper et al., 2024). While the lack of a significant N400-like results in the 9-10-year-old group is similar to the lack of N400-like results in adult-like learning (object-label-ordering), the 9-10-year-olds did indicate above chance learning for the behavioral measures while the label-object-ordering adult group did not (cf. Cosper et al., 2022, 2024), despite comparable accuracy results of 54% for adult group and 57.5% for 9-10-year-olds (cf. Cosper et al., 2022). Furthermore, the lack of N400-like results for the testing phase for the 5-6-year-olds did not parallel findings for the infants (Cosper et al., 2020). As such, the first hypothesis was not confirmed as neither group elicited an N400-like response to violations of sound-pseudoword pairings. Furthermore, our second hypothesis was not confirmed. Younger children did not show earlier and larger N400-like effects or larger behavioral effects than 9-10-year-olds. These results suggest that the shift from auditory dominance in early childhood to the visual dominance after the age of 8 years did not correlate to the shift between infant-like and adult-like auditory associative word learning, as proposed by Cosper and colleagues (2020, 2022, 2024) and Cosper (2020).

One consideration that may have affected associative word learning in both younger and older children is the task itself. While in EEG studies adult learning is modulated by active-learning paradigms (Breitenstein et al., 2005), the effect of active versus passive auditory word learning paradigms is unknown for younger children. Future studies should investigate this in a within design to determine any effects of behavioral measurement on word learning outcome. Another consideration for the lack of N400-like results may be age and cortical development. In the development of the human brain, infants and young children have a higher density of neurons, with 2-year-olds having neuronal density at 55% higher than adults (Huttenlocher, 1979). Between these neurons, cell to cell connections, or synapses, allow for controlled transfer of electrical or chemical signals (cf. Petzoldt & Sigrist, 2014). Huttenlocher (1979) described the synaptic density of these connections in the middle frontal gyrus of 6-month-olds to 7-year-olds as being significantly higher than adult synaptic density. Furthermore, the synaptic density of various brain regions reaches its maximum at different times in development, for example 4 months for the visual cortex or 4 years for the prefrontal cortex (Lenroot & Giedd, 2006). Higher synaptic density has been observed to result in higher capacities of learning (cf. Rakic et al., 1994). As such, it could be that the window of heightened learning for auditory associative word learning has already passed with the 5-6-year-old participants in the current study, but was still a factor in the successful learning and broad distribution of the N400-like effect for the 10-12-month-old infants (Cosper et al., 2020). This interpretation would not only account for the results of the current study, but also allow for the postulation that both younger and older children might be sensitive to modulations to the auditory associative word learning paradigm in a similar manner to adults (object modality, synchrony, and saliency of stimulus; cf. Cosper et al., 2022, 2024). Future studies should focus on 2-4-year-old children in order to explore the possible influence of synaptic density as well as explore stimulus-related and presentation-related task manipulations across development (cf. Cosper et al., 2022, 2024).

### 4.2 Episodic Short-Term Memory Recognition

Episodic short-term memory recognition was tested in three separate STM tasks: auditory-nonverbal, visual-nonverbal, and auditory-verbal. For the younger and older children, the exploratory analysis of all three of the STM tasks yielded STM recognition performances significantly above chance. The strongest measure of STM for each group was in the visual-nonverbal task. While younger children performed quite similarly in the both auditory tasks, older children showed marginally higher recognition memory for auditory-verbal stimuli than auditory-nonverbal stimuli. In all three measures, the older children outperformed the younger children.

The general aptitude for visual memory in both age groups mirrored that of STM effects found in stronger retention of visual events than for auditory events in young adults (Bigelow & Poremba, 2014; Cohen et al., 2009; Talamini et al., 2022; Yuval-Greenberg & Deouell, 2007, 2009). For the auditory modality, the older children retained a marginally higher amount of information for auditory-verbal stimuli than auditory-nonverbal stimuli, as suggested by Talamini and colleagues (2022) for young adults. However, for the younger children this was not the case. The results between auditory-verbal and auditory-nonverbal STM were quite similar, which mirrored results found in statistical learning in young children with auditory-verbal and auditory-nonverbal stimuli (Saffran, 2002; Shufaniya & Arnon, 2018). More important is the development of recognition memory despite stimulus modality. In all three tasks, the older children significantly outperformed the younger children, which conforms to the development of both visual and auditory statistical learning (Raviv & Arnon, 2018; Shufaniya & Arnon, 2018).

In the exploratory correlation analyses, higher auditory-verbal STM performance was significantly correlated to non-musically trained children. This was interesting given the general benefits of musical training in both children and in adults for memory (Chan et al., 1998; Cohen et al., 2011; George & Coch, 2011; Ho et al., 2003) and visual word learning (Dittinger et al., 2017, 2019). This could be an artifact of how musical training was defined in this study, as we defined musical training for the 5-6-year-old age group (minimum of 1.5 years of training) and applied this minimum criterion to the 9-10-year-olds as well. Furthermore, musical training was not a significant factor in higher overall hit-rates, which underscores this possibility. In Dittinger and colleagues (2017), musical training was much more rigidly defined with participants with an average age of 11 years and average musical training of 4.9 years. Had our inclusion criteria for the musical training group been more rigid, we may have also seen different results. This should be considered and directly tested in future studies.

When comparing recognition memory to auditory associative word learning, only visual-nonverbal recognition memory played a significant role. Higher visual-nonverbal recognition memory was positively correlated to higher learning scores in the auditory associative word learning task, despite age. One possible interpretation of this is the general stronger visual memory and statistical learning in both children and adults (Bigelow & Poremba, 2014; Cohen et al., 2009; Raviv & Arnon, 2018; Talamini et al., 2022; Yuval-Greenberg & Deouell, 2007, 2009). Another possibility, yet not mutually exclusive, is the notion that auditory imagery (maintenance of auditory information in a mechanism other than the phonological loop) may interfere with auditory perception and thus with the building of auditory memory during the word learning task (for a review, see Hubbard, 2010). Specifically, interference may occur when processing capacity or resources are exceeded, when auditory imagery coincides with the auditory stimulus, or auditory imagery interferes with auditory detection (Hubbard, 2010). However, this is impossible to disentangle in the current study, as we did not manipulate STM maintenance nor did we analyze any brain signal that correlates to memory encoding or maintenance. As such, the hypothesis put forth by Cosper and colleagues (2022, 2024) was not directly tested in this study and cannot be confirmed or rejected. Future studies should address the neural oscillations of associative word learning, specifically analyzing the theta frequency, which has been shown to correlate to memory encoding and maintenance (Clarke et al., 2018; Hsieh & Ranganath, 2014; Klimesch et al., 1997; Köster & Gruber, 2022; Mölle et al., 2002; White et al., 2013). Although outside the scope of the current study, the neural oscillations should be analyzed in order to attempt disentanglement and to assess further details of the relationship between memory encoding and memory maintenance on associative word learning processes.

### 4.3 Language Development

Language development, specifically receptive vocabulary size, has been previously shown to influence the N400-like response in associative word learning in infants (Junge et al., 2021). As such, we aimed to explore the relationship between language development and auditory associative word learning in 5-6- and 9-10-year-olds. Our exploratory analysis suggested that the cumulative language development score from the SET test battery (Petermann, 2012; Rißling et al., 2013) was a significant factor in higher corrected hit-rates (including all STM and AAWL measures). However, the cumulative SET score did not significantly correlate to the specific hit-rates of any of the STM or AAWL scores. The analysis did yield significant positive correlations between the cumulative score and group or age, which was to be expected over the course of development. Furthermore, the cumulative SET score was significantly higher for female participants than for male participants. While sex differences in language development are not uncommon, many other factors must also be considered with sex, such as environmental factors, age, and task differences (for reviews, see Etchell et al., 2018; Rinaldi et al., 2023). In our exploratory correlation analysis, sex was not correlated with any other measure beyond the language development scores. We interpreted this as sex being a compound factor, which would need to be investigated further, as sex differences in language development overall yield smaller effects than once believed and are not consistent throughout various tasks (Etchell et al., 2018; Rinaldi et al., 2023).

Our findings demonstrated that language continues to develop between the ages of 5 and 10 and is in line with the normalization of the total score as well as the scores of each individual subsections of language development tested in the SET 5-10 (Petermann, 2012). The increases in specific language development areas include vocabulary, semantic relation/categories, (visual) processing speed, language comprehension, language production, and morphology/syntax, which not only improve with age, but higher scores were correlated with the group level and with female participants. One of the subsections, auditory abilities (spoken pseudoword repetition), was not correlated with age or group, but higher scores were correlated to female participants. This is not surprising, as Petermann (2012) developed this assessment only for younger participants, ages 5-6 years. Accordingly, this particular language assessment area was designed to identify phonological working memory delays or deviations from the normalized scores in younger children (Petermann, 2012). Given that the younger and older participants did not significantly differ in this area, our assumption is that this particular area of language development has reached a ceiling effect, or at least a plateau, and is in line with normalized development. Overall, language development had an effect on the overall STM and AAWL corrected hit-rate scores, but did not correlate to higher scores in any specific test. Given that there were no significant N400-like effects in the testing phase for either age group or subgroup, we were unable to explore how language development affected the N400, as has been shown in infant word learning studies (for a review, see Junge et al., 2021).

### 4.4 Limitations and Outlook

In the current study, we were able to assess many aspects of auditory associative word learning in 5-6-and 9-10-year-olds with respect to STM and language development. However, we caution against overgeneralizations of the presented results. One of our limitations in the current study is the AAWL task in three aspects – object modality comparisons, temporal synchrony of stimulus presentation, and task design. While previous investigations into associative word learning with young adults compared the auditory-object and visual-object associative word learning directly against each other (cf. Cosper et al., 2022), this was not done in this study. As such, our interpretations and implications are limited to the scope of auditory associative word learning and may not apply to associative word learning in general.

Furthermore, the AAWL task in the current study only looked at sequential presentation of the environmental sounds and pseudowords, as infants were able to learn these pairings (Cosper et al., 2020) while adults showed difficulty (Cosper et al., 2022, 2024). However, Cosper and colleagues (2022) additionally assessed the factor of stimulus synchrony, testing sequential learning with a 600 ms gap between stimuli and simultaneous learning with a 500 ms overlap of stimuli. While adults were unable to reliably map pseudowords onto environmental sounds in the sequential presentation, they were able to show above chance learning in both behavioral measures and with N400-like effects in the testing phase with simultaneous presentation (Cosper et al., 2022). However, these two different temporal synchronies were not directly tested in the current study, as we solely focused on the sequential presentation. Finally, the AAWL task in the current study was presented as an active learning task, meaning that behavioral measures were also collected in the testing phase. While infants were able to show learning effects in a passive auditory associative word learning design (Cosper et al., 2020), active tasks facilitate adult learning (Breitenstein et al., 2005). It is unknown to what extent active or passive designs have an influence on auditory associative word learning in children. Thus, future research into auditory associative word learning should attend to these limitations by experimentally considering modality comparisons, temporal comparisons, and task design to deepen our understanding of learning factors for associative word learning in children.

For the current study, we focused on exploring a possible age when children shift from infant-like learning to more adult-like learning. For this, we highlighted the hypothesis that younger children exhibit an auditory dominance while older children shift to a visual dominance, this hypothesis further expanded to state that formal education could also have an effect on modality dominance (Nava & Pavani, 2013). In our sample of 5-6-year-olds, not all of the children were tested before entering into the formal education system. As such, we were unable to assess the claim that schooling may have an effect on auditory associative word learning. Another age-related limitation is the choice of the age groups in the current study. We argued that there may be a shift from infant-like auditory associative word learning to adult-like learning along the same lines as shifts in modal dominance and emerging effects of multisensory integration (Gori et al., 2008; Nardini et al., 2008; Nava & Pavani, 2013); however, we were unable to find evidence for this in the current study. As such, we can reject the supposition that modality dominance has a general effect on auditory associative word learning. Instead, we have suggested that successful infant-like learning (cf. Cosper et al., 2020) may result from high synaptic density in synaptogenesis during the early years of life, which may no longer factor in with the selected age groups in the current study (Huttenlocher, 1979; Lenroot & Giedd, 2006; Rakic et al., 1994).

With regard to STM, we also caution overgeneralization of the current results. Our episodic STM recognition tasks were presented in isolation, only assessing recognition memory. In our measurement of STM, we did not test the visual-verbal modality, as children in Germany do not necessarily learn to read before entering into the formal education system. Without direct testing, we cannot explore how the development of visual-verbal recognition might affect associative word learning in any modality. Furthermore, the current study focused solely on recognition memory. In a meta-analysis of younger and older adults, the measurement of memory (recognition vs. recall) indicated differing effects with regard to age, where age differences in recognition memory were smaller than for free recall (Rhodes et al., 2019). While recognition memory tested in the current study yielded interesting results, free recall should also be considered in the future to understand how memory affects associative word learning on a broader level. Furthermore, direct testing of memory maintenance processes was not conducted in the current set of experiments. As such, we are unable to answer the question if specific auditory maintenance processes are being interrupted or at least affected by the presentation and order of auditory-verbal and auditory-nonverbal stimuli (Baddeley, 2012; Hubbard, 2010; Li et al., 2024). For this, further investigation is necessary with targeted manipulations of memory maintenance.

Finally, we acknowledge the limitations within our exploratory analyses of sex and musical training. While a few significant correlations were found between sex and language development, it is important to note that these effects are not consistent across age or task and are not consistent with regard to the development of the brain (Etchell et al., 2018; Rinaldi et al., 2023). Sex differences were present in the current study, but we do not suggest that sex facilitates or inhibits language development or auditory associative word learning outcomes in children. Lastly, musical training has been shown to facilitate associative word learning in the auditory modality of young adults (Cosper et al., 2024) as well as across the lifespan in the visual modality (Dittinger et al., 2019). However, these studies had clear and age appropriate categorization criteria for musicianship and non-musicianship for the purpose of direct comparison. The current study loosely defined musical training for the 5-6-year-old age group and applied the same criterion for the 9-10-year-olds. Further investigation is needed to fully understand the impact of musical training on auditory associative word learning in both of the age groups in the current study, with clear and age-appropriate inclusion criteria and balanced sample sizes.

## 5 Conclusion

In the current study, we were able to show that auditory associative word learning is similarly difficult with 5-6- and 9-10-year-olds as it is with young adults. Furthermore, higher visual-nonverbal memory retention was positively correlated with higher auditory associative word learning measurements. While language development played a general role in higher overall behavioral measures of learning, it was not correlated to any specific STM or associative word learning task. Musical training affected the N400-like response of the training phase in auditory associative word learning in older children; yet, did not correlate to any higher behavioral measures in associative word learning or STM. Overall, we were able to show that sequential auditory associative word learning is a difficult task, which may be modulated by factors of age, musical training, and visual STM. We further postulated influences of object modality, temporal synchrony of stimulus presentation, memory maintenance, task design, and synaptogenesis on associative word learning, indicating their importance for future studies.

## Data Availability Statement

The data and scripts used for the analysis can be found here: https://osf.io/xmr5q/ and the experiments were not preregistered.

## Declaration of generative AI and AI-assisted technologies in the manuscript preparation process

The authors declare that no generative AI or AI-assisted technologies were used in the writing of the analysis or the creation of figures and tables in the manuscript or in the supplementary materials.

## Author contributions

**Samuel H. Cosper:** Conceptualization, Data Curation, Formal analysis, Investigation, Funding acquisition, Project administration, Visualization, Writing – original draft, Writing – review and editing. **Laura Bachmann:** Formal analysis, Investigation, Visualization, Writing – original draft, Writing – review and editing. **Eleonora Sehmer:** Investigation, Writing – review and editing. **Alexandra Steidel:** Investigation, Writing – review and editing. **Shu-Chen Li:** Resources, Supervision, Writing – review and editing.

## Acknowledgements

The authors would like to thank Anna Weinert and Charline Lucie Kyra Friebel for assisting with data collection and their insights on the data processing. We would also like to thank James Bigelow and Michael Cohen for their interest and for answering questions about short-term episodic recognition memory, and Brais Gonzalez Sousa and Heidrun Schultz for their assistance with the R scripts and Xue-Rui Peng for her assistance with the Matlab figure scripts. And, of course, we could not have presented these results without all of the participants and their families – thank you.

