## Supplementary Materials for "The Interplay between Language Development, Short-Term Memory, and Auditory Associative Word Learning in Younger and Older Children"

### Supplementary Materials (A-K)

#### A. List of pseudowords used in the AAWL task (listed alphabetically):

1) Diebust

2) Dirfel

3) Dorten

4) Imsten

5) Knaugel

6) Krissel

7) Maffau

8) Mofter

9) Mottich

10) Niller

11) Pipfen

12) Saden

13) Tolzen

14) Wotir

15) Zorgel

16) Zufel

#### B. Alphabetical list of environmental sounds used in the AAWL task (taken from the NESSTI database for environmental sounds (Hocking et al., 2013)):

1) Canary bird tweet

2) Coffee machine

3) Coin spin

4) Computer boot

5) Cricket chirp

6) Donkey sound

7) Fire burning

8) Fly buzzing

9) Gong ringing

10) Jingling keys

11) Knife chopping

12) Maracas shaking

13) Saxophone

14) Water bubbling

15) Whale call

16) Windchimes

#### C: List of stimuli for the STM auditory-nonverbal task:

| Number | Sound | Source |
|---|---|---|
| 1 | Church bell | Gygi & Shafiro, 2010 |
| 2 | Car start | Gygi & Shafiro, 2010 |
| 3 | glass smash | Gygi & Shafiro, 2010 |
| 4 | ice drop into glass | Gygi & Shafiro, 2010 |
| 5 | shovel - dig | Gygi & Shafiro, 2010 |
| 6 | bowling - strike | Gygi & Shafiro, 2010 |
| 7 | pingpong | Gygi & Shafiro, 2010 |
| 8 | toilet flush | Gygi & Shafiro, 2010 |
| 9 | airplane | Gygi & Shafiro, 2010 |
| 10 | basketball | Gygi & Shafiro, 2010 |
| 11 | bubbles | Gygi & Shafiro, 2010 |
| 12 | buzzer | Gygi & Shafiro, 2010 |
| 13 | camera | Gygi & Shafiro, 2010 |
| 14 | car tires (breaks) | Gygi & Shafiro, 2010 |
| 15 | clapping | Gygi & Shafiro, 2010 |
| 16 | clock | Gygi & Shafiro, 2010 |
| 17 | helicopter | Gygi & Shafiro, 2010 |
| 18 | trashbag | Gygi & Shafiro, 2010 |
| 19 | cutting paper | Gygi & Shafiro, 2010 |
| 20 | cymbols | Gygi & Shafiro, 2010 |
| 21 | door | Gygi & Shafiro, 2010 |
| 22 | electric saw | Gygi & Shafiro, 2010 |
| 23 | match strike | Gygi & Shafiro, 2010 |
| 24 | hammer | Gygi & Shafiro, 2010 |
| 25 | horse (run) | Gygi & Shafiro, 2010 |
| 26 | typing | Gygi & Shafiro, 2010 |
| 27 | phone ring | Gygi & Shafiro, 2010 |
| 28 | pour water | Gygi & Shafiro, 2010 |
| 29 | printer | Gygi & Shafiro, 2010 |
| 30 | film projector | Gygi & Shafiro, 2010 |
| 31 | rain | Gygi & Shafiro, 2010 |
| 32 | rocking chair | Gygi & Shafiro, 2010 |
| 33 | screen (door) | Gygi & Shafiro, 2010 |
| 34 | stapler | Gygi & Shafiro, 2010 |

| | | |
|---|---|---|
| 35 | thunder | Gygi & Shafiro, 2010 |
| 36 | train | Norman-Haignere et al., 2015 |
| 37 | chewing | Norman-Haignere et al., 2015 |
| 38 | whistle | Gygi & Shafiro, 2010 |
| 39 | windshield wipers | Gygi & Shafiro, 2010 |
| 40 | zipper | Gygi & Shafiro, 2010 |
| 41 | baby cry | Gygi & Shafiro, 2010 |
| 42 | cat (meow) | Gygi & Shafiro, 2010 |
| 43 | monkey (chimp) | Gygi & Shafiro, 2010 |
| 44 | cough | Gygi & Shafiro, 2010 |
| 45 | Cow (moo) | Gygi & Shafiro, 2010 |
| 46 | dog (bark) | Gygi & Shafiro, 2010 |
| 47 | drums | Gygi & Shafiro, 2010 |
| 48 | electric guitar | Gygi & Shafiro, 2010 |
| 49 | flute | Gygi & Shafiro, 2010 |
| 50 | footsteps | Gygi & Shafiro, 2010 |
| 51 | gargling (human) | Gygi & Shafiro, 2010 |
| 52 | harp | Gygi & Shafiro, 2010 |
| 53 | laugh (human) | Gygi & Shafiro, 2010 |
| 54 | horse (neigh) | Gygi & Shafiro, 2010 |
| 55 | sheep (bah) | Gygi & Shafiro, 2010 |
| 56 | police siren | Gygi & Shafiro, 2010 |
| 57 | sneeze (human) | Gygi & Shafiro, 2010 |
| 58 | cash register | Gygi & Shafiro, 2010 |
| 59 | rooster | Gygi & Shafiro, 2010 |
| 60 | bird call | Gygi & Shafiro, 2010 |
| 61 | alarm clock | Norman-Haignere et al., 2015 |
| 62 | bike bell | Norman-Haignere et al., 2015 |
| 63 | car horn | Norman-Haignere et al., 2015 |
| 64 | coin in vending machine | Norman-Haignere et al., 2015 |
| 65 | duck (quack) | Norman-Haignere et al., 2015 |
| 66 | flying (wings flapping) | Norman-Haignere et al., 2015 |
| 67 | frog | Norman-Haignere et al., 2015 |
| 68 | frying | Norman-Haignere et al., 2015 |
| 69 | grunting (human) | Norman-Haignere et al., 2015 |
| 70 | hair dryer | Norman-Haignere et al., 2015 |

| | | |
|---|---|---|
| 71 | microwave button | Norman-Haignere et al., 2015 |
| 72 | page turn (newspaper) | Norman-Haignere et al., 2015 |
| 73 | shuffling cards | Norman-Haignere et al., 2015 |
| 74 | spray can | Norman-Haignere et al., 2015 |
| 75 | toothbrush | Norman-Haignere et al., 2015 |
| 76 | vacuum | Norman-Haignere et al., 2015 |
| 77 | squeaky toy | Norman-Haignere et al., 2015 |
| 78 | computer start (windows) | Norman-Haignere et al., 2015 |
| 79 | writing on paper | Norman-Haignere et al., 2015 |
| 80 | truck backing up beep | Norman-Haignere et al., 2015 |
| 81 | tv/radio static | Norman-Haignere et al., 2015 |
| 82 | motorcycle engine | Norman-Haignere et al., 2015 |
| 83 | water kettle | Norman-Haignere et al., 2015 |
| 84 | dishes | Norman-Haignere et al., 2015 |
| 85 | bees | Norman-Haignere et al., 2015 |
| 86 | breathing (human) | Norman-Haignere et al., 2015 |
| 87 | cell phone vibrating | Norman-Haignere et al., 2015 |
| 88 | chair rollinig | Norman-Haignere et al., 2015 |
| 89 | dog (drinking/slurping) | Norman-Haignere et al., 2015 |
| 90 | electric shaver | Norman-Haignere et al., 2015 |
| 91 | whistling (human) | Norman-Haignere et al., 2015 |
| 92 | busy signal (beep) | Norman-Haignere et al., 2015 |
| 93 | iPhone ringtone | Norman-Haignere et al., 2015 |
| 94 | heart beat | Norman-Haignere et al., 2015 |
| 95 | waving metal sheet | freesound.org |
| 96 | opening soda can | freesound.org |

#### D: List of stimuli for the STM auditory-verbal task (English translations in a separate list below):

**English translations of spoken words for the auditory-verbal STM task (for German list, please contact first/corresponding author:**

| Number | Noun | A)djective | Adverb/ prep/ pronoun | Verb |
|---|---|---|---|---|
| 1 | Life | Known | Between | To change |
| 2 | August | Total | Enough | To demand |
| 3 | Member | Slow | Anyway | To help |
| 4 | Daughter | Constant | Primarily | To smile |
| 5 | Business | Normal | Nevertheless | To open |
| 6 | Border | Private | Currently | To jump |
| 7 | Newspaper | Central | Mostly | To buy |
| 8 | Technology | Remaining | In front | To build |
| 9 | Body | Calm | Outside | To realize |
| 10 | Culture | Cheap | There | To love |
| 11 | Opinion | Half | To date | To turn |
| 12 | Summer | Yellow | After | To rustle |
| 13 | Future | Typical | Over | To party |
| 14 | Order | Ready | Never | To follow |
| 15 | Direction | Quiet | Indeed | To be sickly |
| 16 | Attempt | Fierce | Finally | To force |
| 17 | Region | Clerical | Without | To calculate |
| 18 | Food | Critical | Excessively | To protect |
| 19 | Car | Huge | Back | To achieve |
| 20 | Tip | West | Twice | To dream |
| 21 | Environment | Kind | Midway | To fill |
| 22 | Household | Clear | Over there | To measure |
| 23 | Novel | Written | Not a | To plant |
| 24 | Shoulder | Powerful | At most | To learn |

**E: List of stimuli for the STM visual-nonverbal task (taken from the Amsterdam Library of Object Images (Geusebroek et al., 2005)):**

| Number | Item description | item number in database |
|---|---|---|
| 1 | Lemon | 257_l |
| 2 | brass weight | 260_l |
| 3 | leather ball | 266_r |
| 4 | nail brush | 267_r |
| 5 | thread | 275_l |
| 6 | light bulb | 293_c |
| 7 | dumpling press | 294_c |
| 8 | lipstick case | 316_r |
| 9 | shell | 329_l |
| 10 | vase | 332_c |
| 11 | lego duplo toy | 339_c |
| 12 | glass with toothpicks | 351_c |
| 13 | small cloth bag | 353_l |
| 14 | rubber ducky | 354_r |
| 15 | hand statue | 373_c |
| 16 | cat toy | 375_r |
| 17 | birthday blow toy | 391_l |
| 18 | whistle | 393_l |
| 19 | massage toy | 400_r |
| 20 | toy mouse | 403_c |
| 21 | hole punch | 405_r |
| 22 | snowman | 406_c |
| 23 | hour glass | 419_r |
| 24 | dart | 421_c |
| 25 | Kinder egg toy pill | 429_c |
| 26 | green/white tube | 439_l |
| 27 | Book | 442_l |
| 28 | computer mouse | 457_l |
| 29 | pink pig | 459_l |
| 30 | toilet paper roll | 461_c |
| 31 | donut | 479_l |
| 32 | toy car | 482_r |

| | | |
|---|---|---|
| 33 | tape despensor | 488_r |
| 34 | wooden box | 493_l |
| 35 | nose spray | 497_c |
| 36 | green bird | 503_c |
| 37 | coffee cup | 507_c |
| 38 | plant pot | 508_r |
| 39 | water cup | 514_r |
| 40 | broom head | 524_l |
| 41 | top | 553_r |
| 42 | wooden doll | 614_c |
| 43 | electric plug | 622_l |
| 44 | easter egg | 626_r |
| 45 | eye-dropper | 629_r |
| 46 | paperclips in box | 635_c |
| 47 | stapeler | 642_l |
| 48 | Ramubtan | 664_l |
| 49 | bike light | 694_l |
| 50 | clock | 701_l |
| 51 | fish | 731_c |
| 52 | flower | 738_l |
| 53 | clay dish | 748_c |
| 54 | stacking toy | 769_r |
| 55 | woven basket | 780_r |
| 56 | pine cone | 785_c |
| 57 | bread | 801_c |
| 58 | watering can | 809_r |
| 59 | tissue box | 815_r |
| 60 | measuring cup | 819_c |
| 61 | shopping basket | 821_r |
| 62 | candle | 839_c |
| 63 | cookie tin | 849_c |
| 64 | water bottle | 856_r |
| 65 | cactus | 866_r |
| 66 | balloon | 871_r |
| 67 | origami fox | 878_l |
| 68 | garden shears | 891_r |

| | | |
|---|---|---|
| 69 | chair | 982_l |
| 70 | duster | 895_l |
| 71 | chinese yo-yo | 903_c |
| 72 | coffee pot | 911_c |
| 73 | drum | 916_r |
| 74 | metal tool | 919_r |
| 75 | egg carton | 924_c |
| 76 | knife block | 926_l |
| 77 | bananas | 950_r |
| 78 | towels | 973_c |
| 79 | hat | 979_c |
| 80 | sponge roll | 980_l |
| 81 | metal pan | 982_r |
| 82 | shells in glass | 985_c |
| 83 | stuffed sheep | 988_r |
| 84 | rice tub | 991_c |
| 85 | staple remover | 358_r |
| 86 | snow globe | 526_r |
| 87 | sticky post-it notes | 468_l |
| 88 | wooden clogs | 417_l |
| 89 | shoe | 826_c |
| 90 | bowl | 30_l7c3 |
| 91 | chess piece | 33_i120 |
| 92 | rope | 72_r20 |
| 93 | wooden boat | 75_r355 |
| 94 | badmitton shuttle | 155_r70 |
| 95 | rubix cube | 200_i130 |
| 96 | frog | 221_i160 |

#### F. Full Electrode Layout

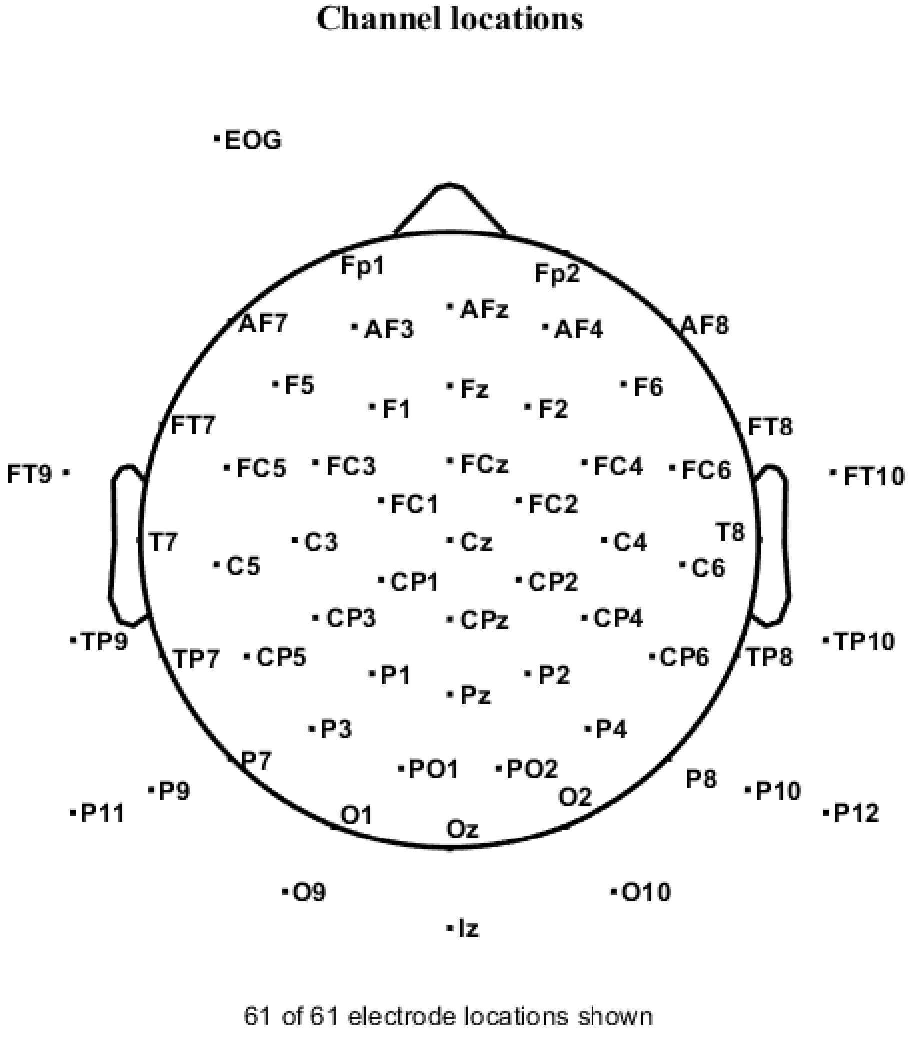


Figure SM_F_1: Approximate electrode location on the scalp. Please note – while the depiction is approximate, electrode site locations are clearly defined in the Matlab scripts and are applied to the data from site locations given by the EEG cap manufacturer. Left/right opposing electrode sites are defined with mirrored locations on the scalp, equidistant from the midline.

#### G. ERP Figures for differences between males and females in the 5-6-year-old age group:

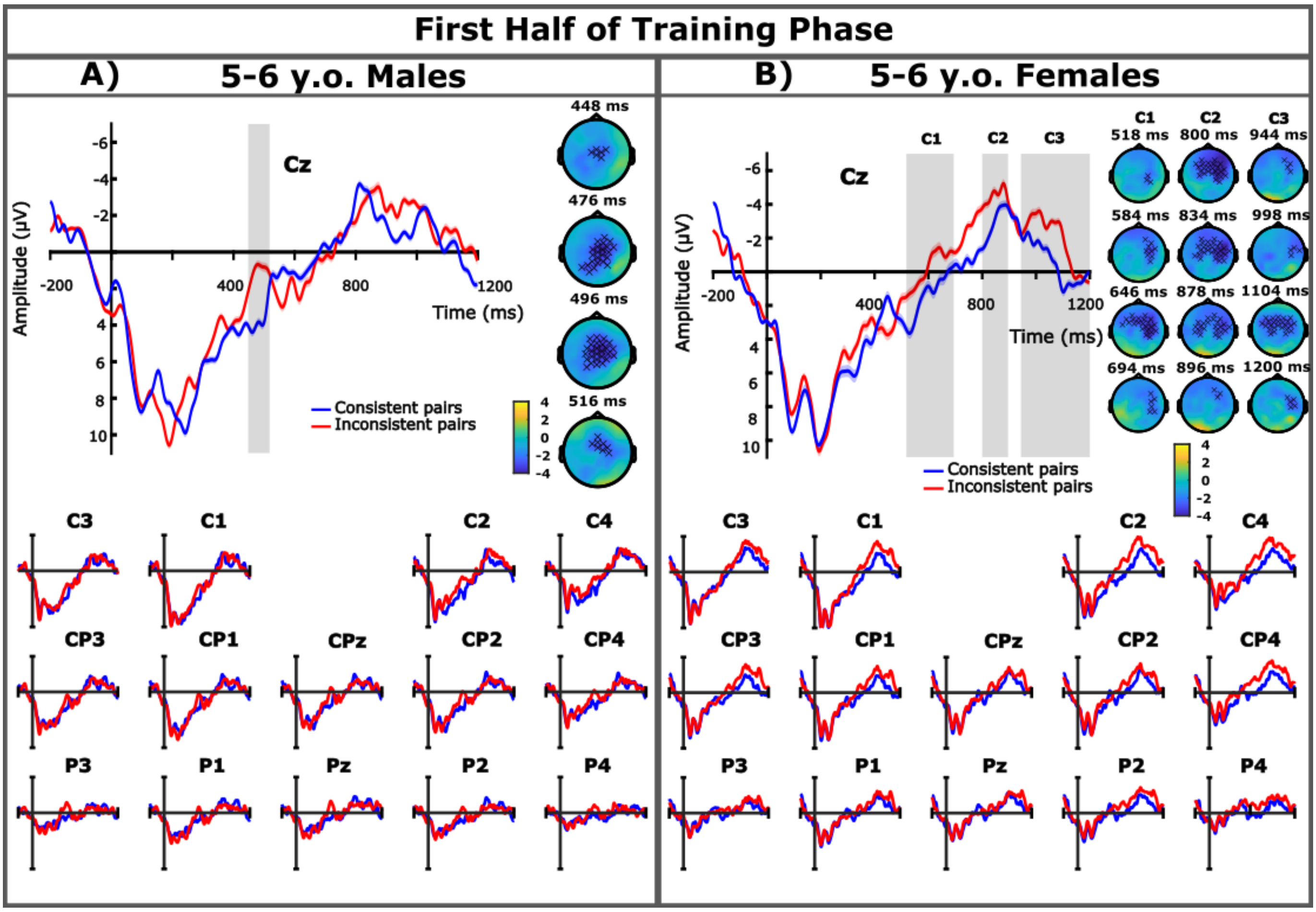


Figure SM_G_1: First half of the training phase results. ERPs are presented time-locked to the onset of the pseudoword for the first half of the training phase of males and females in the 5-6-year-old age group. The Cz electrode was selected as a representative electrode for both groups, further centro-parietal electrodes are displayed below. The blue lines indicate the consistent pairs while the red lines indicate the inconsistent pairs. Gray bar indicates time window in which significant differences between the conditions were found. Topological difference maps from the cluster-based permutation test are presented only where significant differences were found and show the electrode distribution in the cluster over time (the X symbol indicates $p < 0.05$). C1, C2, and C3 in section B refer to the three separate clusters in which significant differences were found.

##### Sex differences: 5-6-year-olds first half of training phase results

For male participants in the first half of training, a negative cluster was found to have significant differences between inconsistent and consistent conditions in the 400 – 800 ms time window between 448 – 518 ms ($p$ = 0.016). No clusters containing significant processing differences was found in the 800 – 1200 ms time window for male participants. See Figure SM_G_1(A). Female participants yielded a cluster of significant processing differences in the 400 – 800 ms time window cluster 1 between 518 – 694 ms ($p$ = 0.034). Furthermore, two additional negative clusters with significant differences were found in the 800 – 1200 ms time window cluster 2 between 800 – 896 ms ($p$ = 0.042) and cluster 3 between 944 – 1200 ms ($p$ = 0.018). See Figure SM_G_1(B).

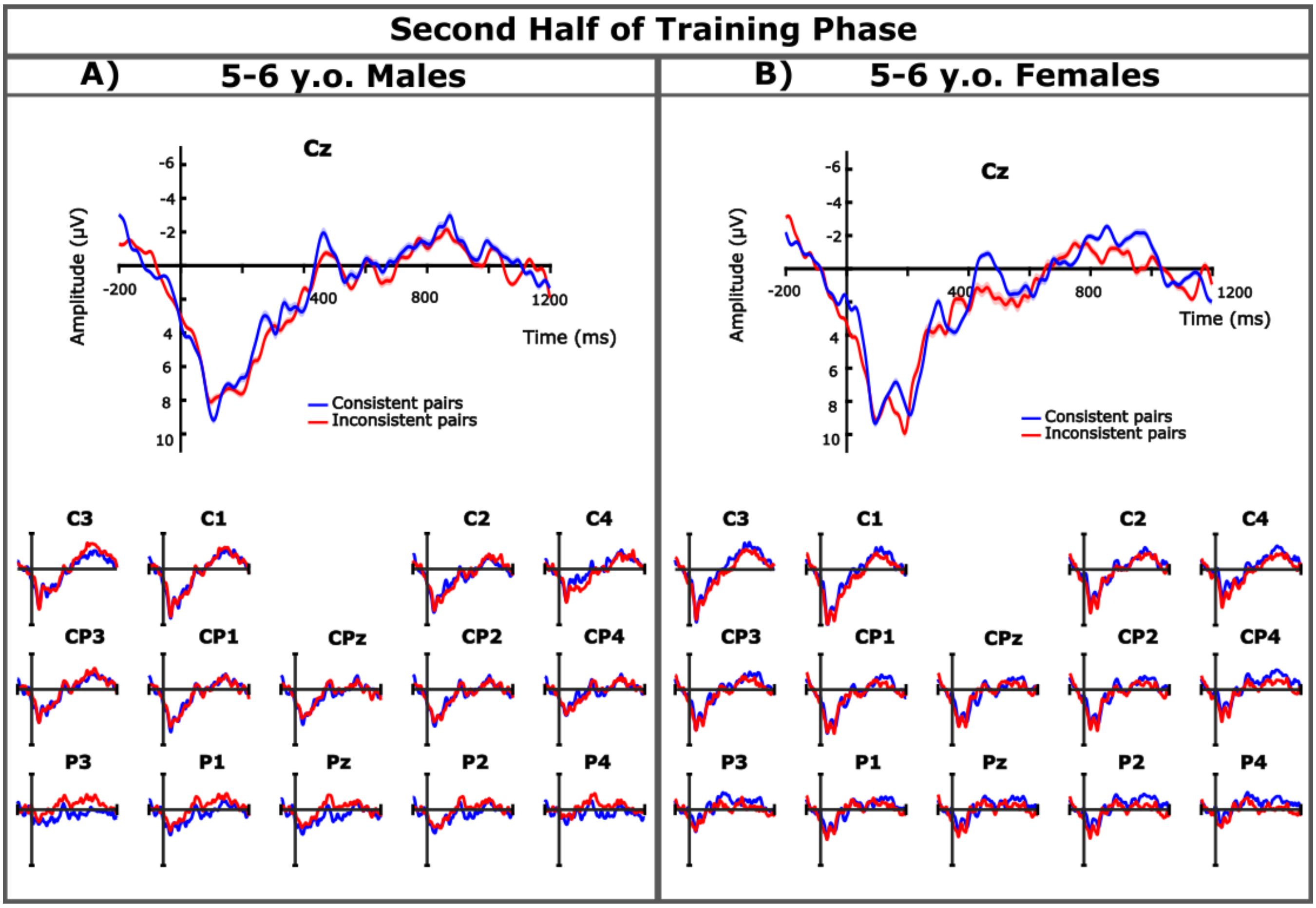


Figure SM_G_2: Second half of the training phase results. ERPs are presented time-locked to the onset of the pseudoword for the second half of the training phase of males and females in the 5-6-year-old age group. The Cz electrode was selected as a representative electrode for both groups, further centro-parietal electrodes are displayed below. The blue lines indicate the consistent pairs while the red lines indicate the inconsistent pairs.

#### Sex differences: 5-6-year-olds second half of training phase results

For the second half of training, no clusters of significant differences were found for male or female participants in either the N400 or later time window (see Figure SM_G_2).

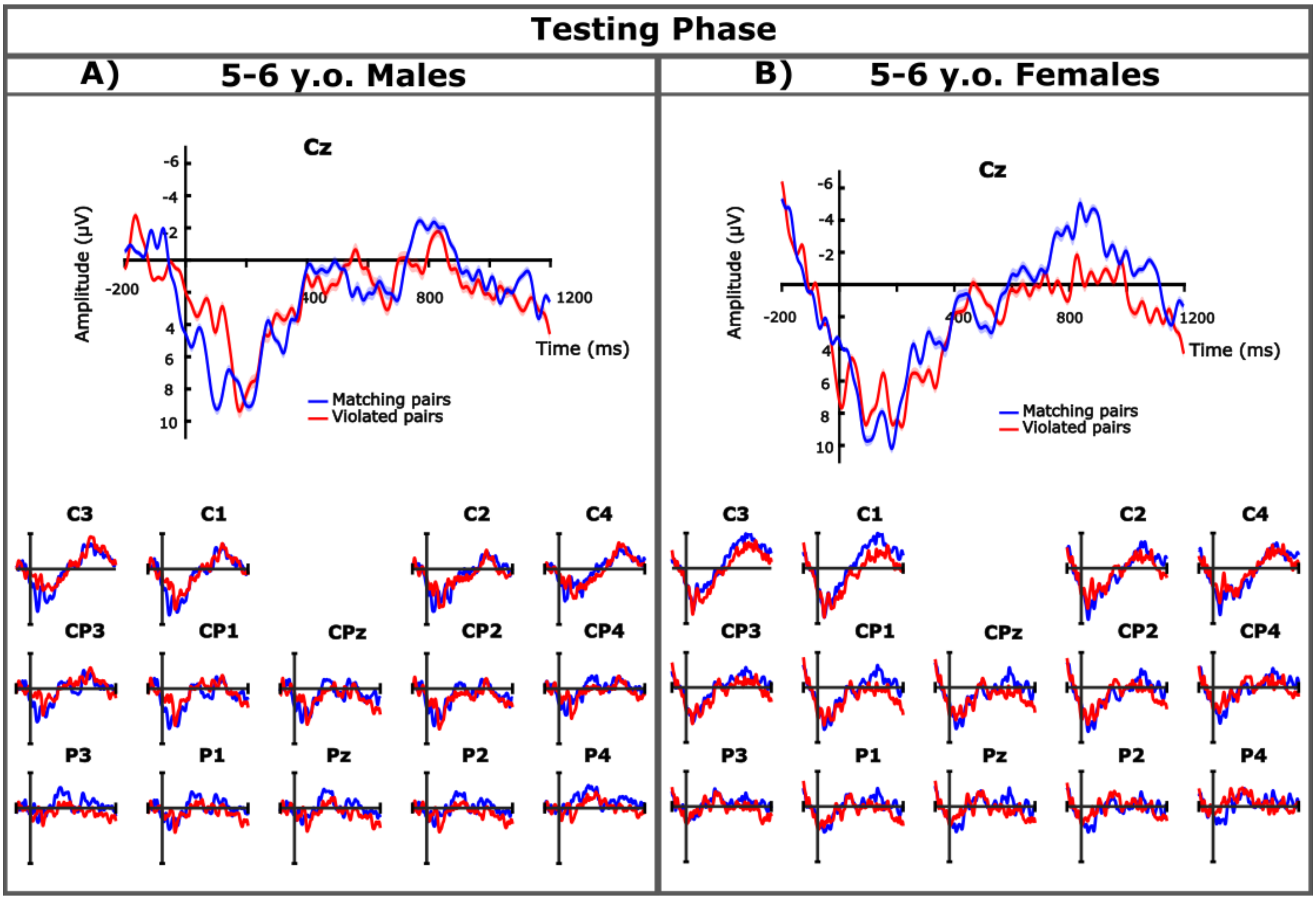


Figure SM_G_3: Testing phase results. ERPs are presented time-locked to the onset of the pseudoword for the testing phase of males and females in the 5-6-year-old age group. The Cz electrode was selected as a representative electrode for both groups, further centro-parietal electrodes are displayed below. The blue lines indicate the matching pairs while the red lines indicate the violated pairs.

#### Sex differences: 5-6-year-olds testing phase results

For the testing phase, no clusters of significant differences were found for male or female participants in either the N400 or later time window (see Figure SM_G_3).

#### H. ERP Figures for differences between males and females in the 9-10-year-old age group:

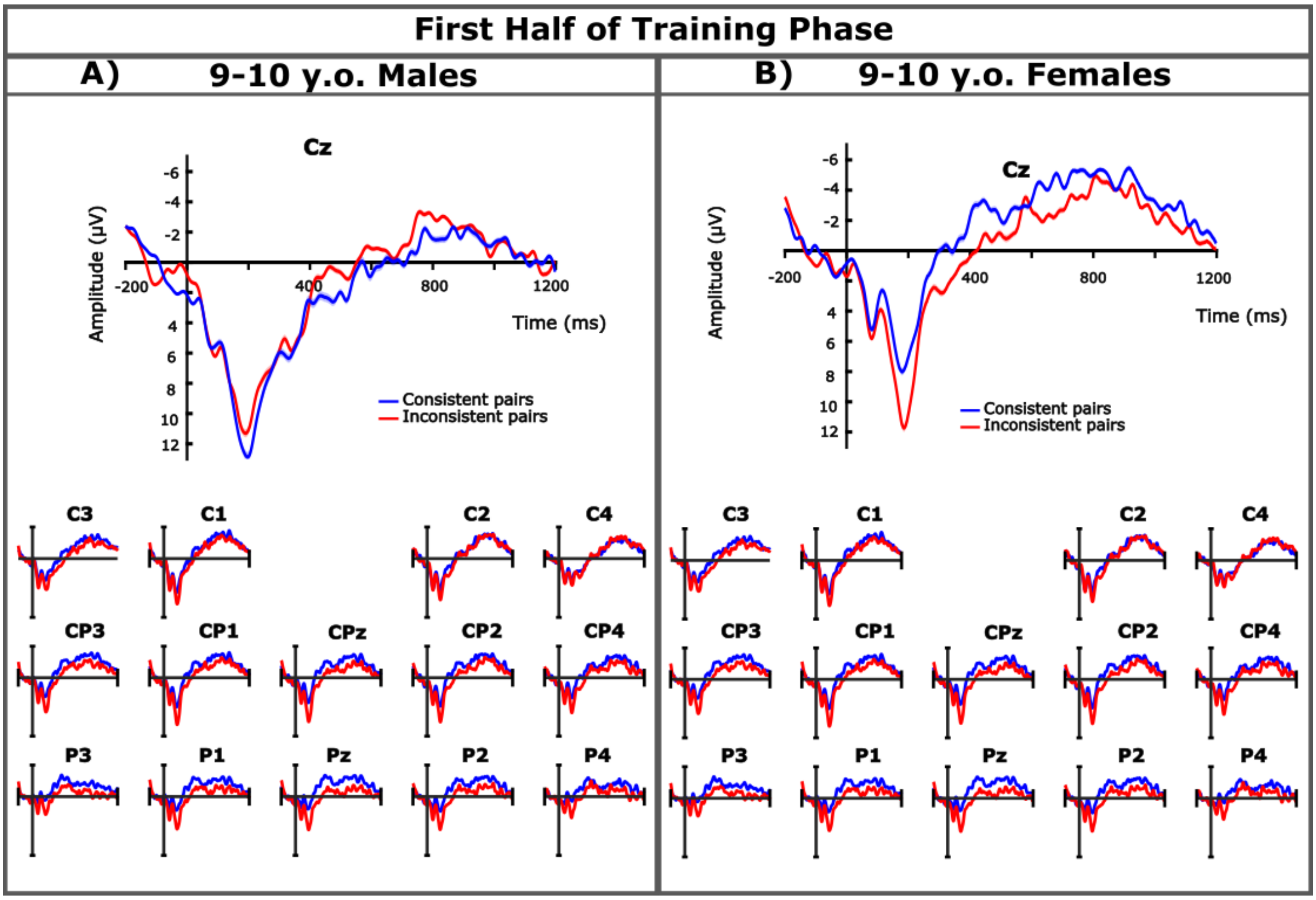


Figure SM_H_1: First half of the training phase results. ERPs are presented time-locked to the onset of the pseudoword for the first half of the training phase of males and females in the 9-10-year-old age group. The Cz electrode was selected as a representative electrode for both groups, further centro-parietal electrodes are displayed below. The blue lines indicate the consistent pairs while the red lines indicate the inconsistent pairs.

##### Sex differences: 9-10-year-olds first half of training phase results

For the first half of training, no clusters of significant differences were found for male or female participants in either the N400 or later time window (see Figure SM_H_1).

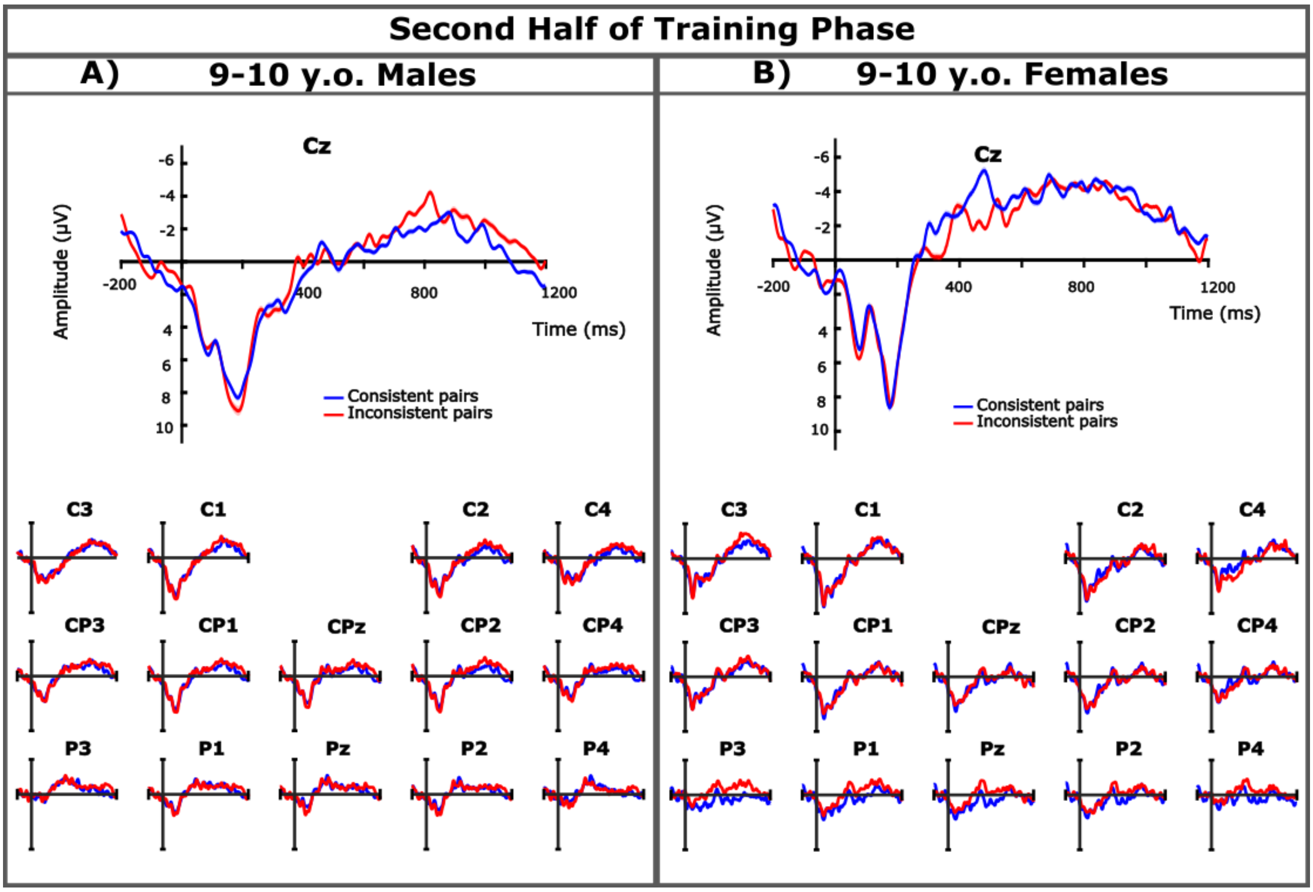


Figure SM_H_2: Second half of the training phase results. ERPs are presented time-locked to the onset of the pseudoword for the second half of the training phase of males and females in the 9-10-year-old age group. The Cz electrode was selected as a representative electrode for both groups, further centro-parietal electrodes are displayed below. The blue lines indicate the consistent pairs while the red lines indicate the inconsistent pairs.

#### Sex differences: 9-10-year-olds second half of training phase results

For the second half of training, no clusters of significant differences were found for male or female participants in either the N400 or later time window (see Figure SM_H_2).

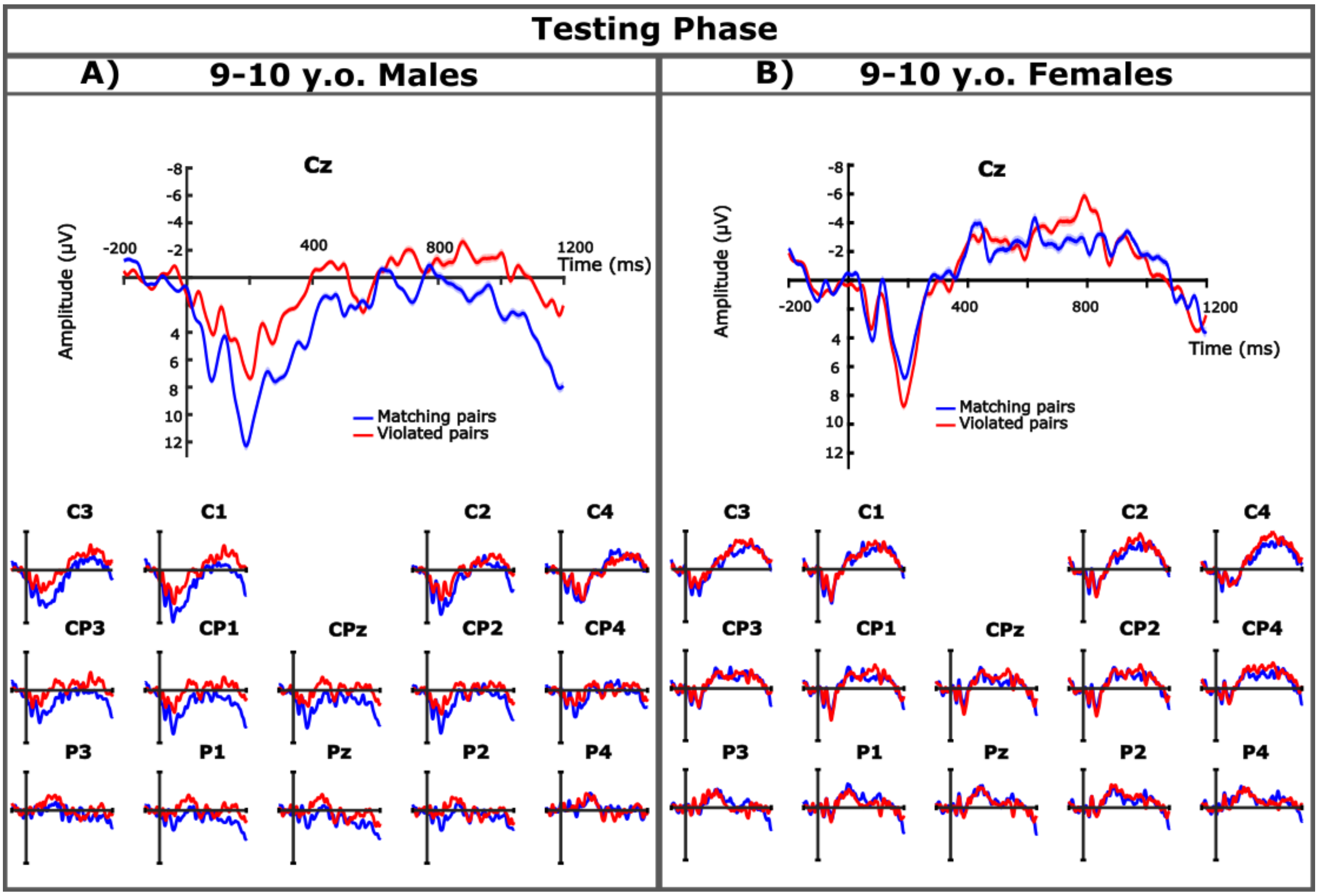


Figure SM_G_3: Testing phase results. ERPs are presented time-locked to the onset of the pseudoword for the testing phase of males and females in the 9-10-year-old age group. The Cz electrode was selected as a representative electrode for both groups, further centro-parietal electrodes are displayed below. The blue lines indicate the matching pairs while the red lines indicate the violated pairs.

#### Sex differences: 9-10-year-olds testing phase results

For the testing phase, no clusters containing significant processing differences were found in the male or female participants in either time window (see Figure SM_H_3).

#### I. ERP Figures for differences between musically-trained and non-musically-trained 9-10 year olds:

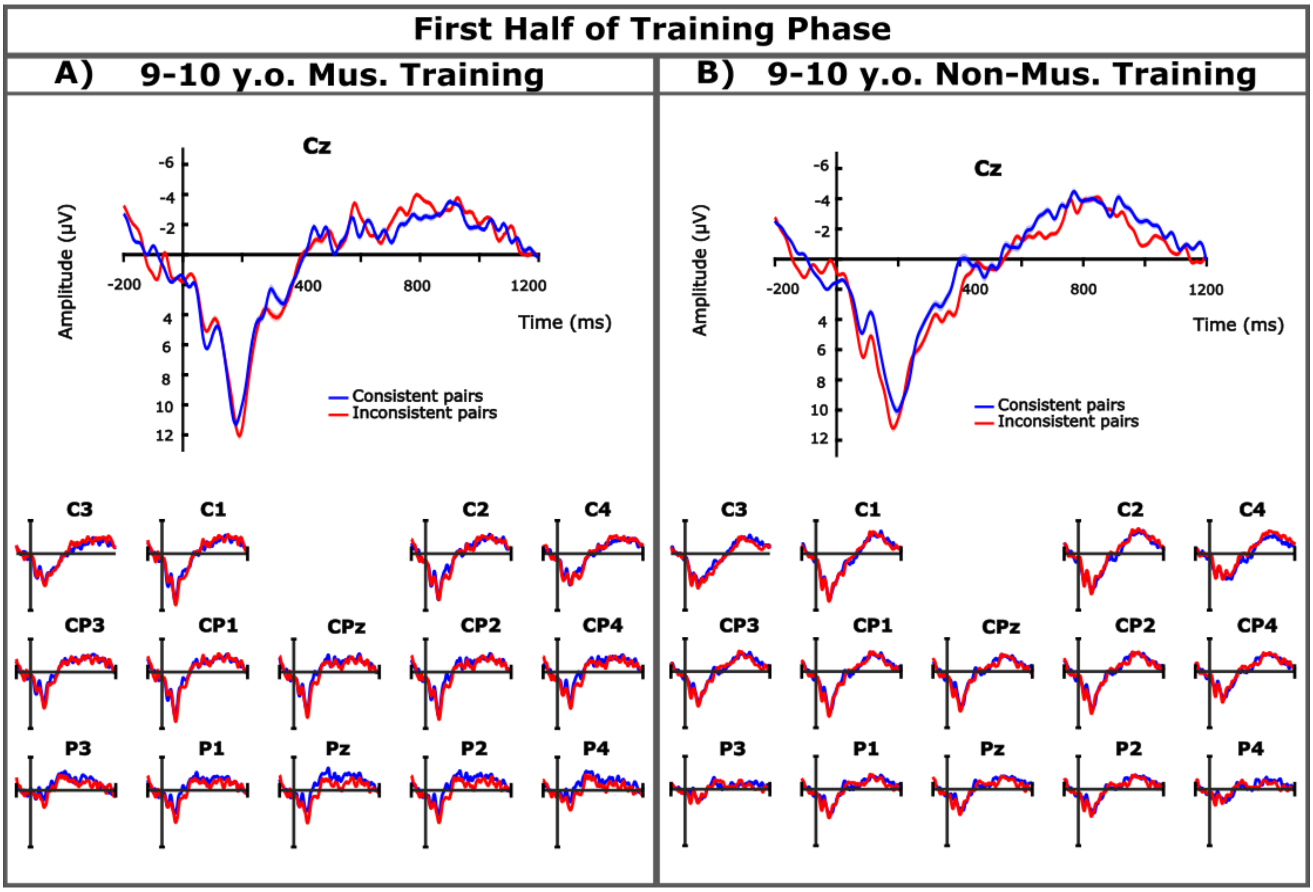


Figure SM_I_1: First half of the training phase results. ERPs are presented time-locked to the onset of the pseudoword for the first half of the training phase of musically-trained and non-musically-trained 9-10 year olds. The Cz electrode was selected as a representative electrode for both groups, further centro-parietal electrodes are displayed below. The blue lines indicate the consistent pairs while the red lines indicate the inconsistent pairs.

##### Musical-training differences: 9-10-year-olds first half of training phase results

No clusters of significant differences were found for male or female participants in either the N400 or later time window (see Figure SM_I_1).

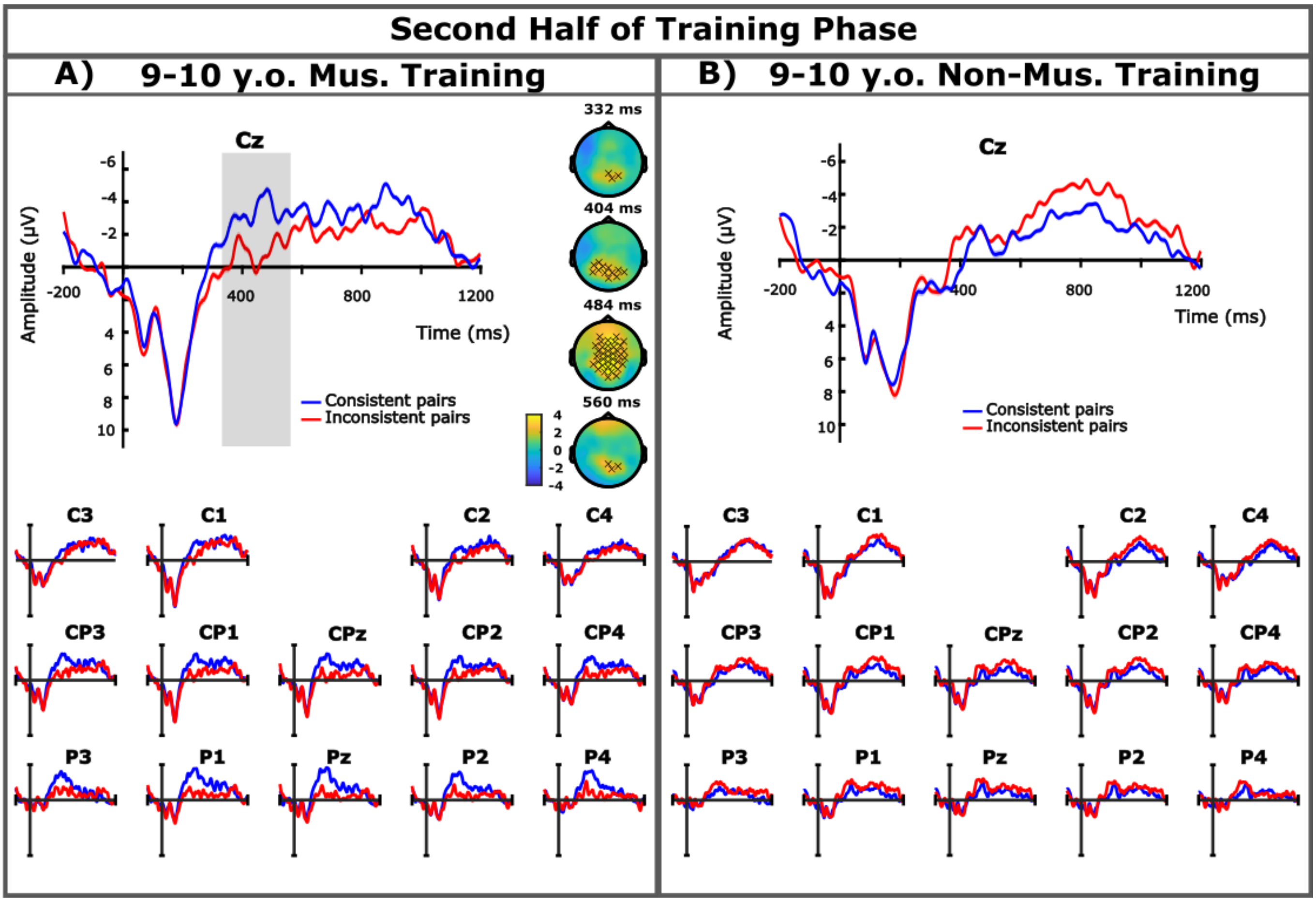


Figure SM_I_2: Second half of the training phase results. ERPs are presented time-locked to the onset of the pseudoword for the second half of the training phase of musically-trained and non-musically-trained 9-10 year olds. The Cz electrode was selected as a representative electrode for both groups, further centro-parietal electrodes are displayed below. The blue lines indicate the consistent pairs while the red lines indicate the inconsistent pairs. Gray bar indicates time window in which significant differences between the conditions were found. Topological difference maps from the cluster-based permutation test are presented only where significant differences were found and show the electrode distribution in the cluster over time (the X symbol indicates p < 0.05).

#### Musical-training differences: 9-10-year-olds second half of training phase results

In the second half of training, musically trained participants yielded a positive cluster with significant processing differences between inconsistent and consistent parings in the N400 time window between 332 – 562 ms (*p* = 0.022), while no clusters with significant differences were found in the later time window (see Figure SM_I_2(A)). Non-musically trained participants also yielded no clusters of significant processing differences in either the N400 or the later time window (see Figure SM_I_2(B))

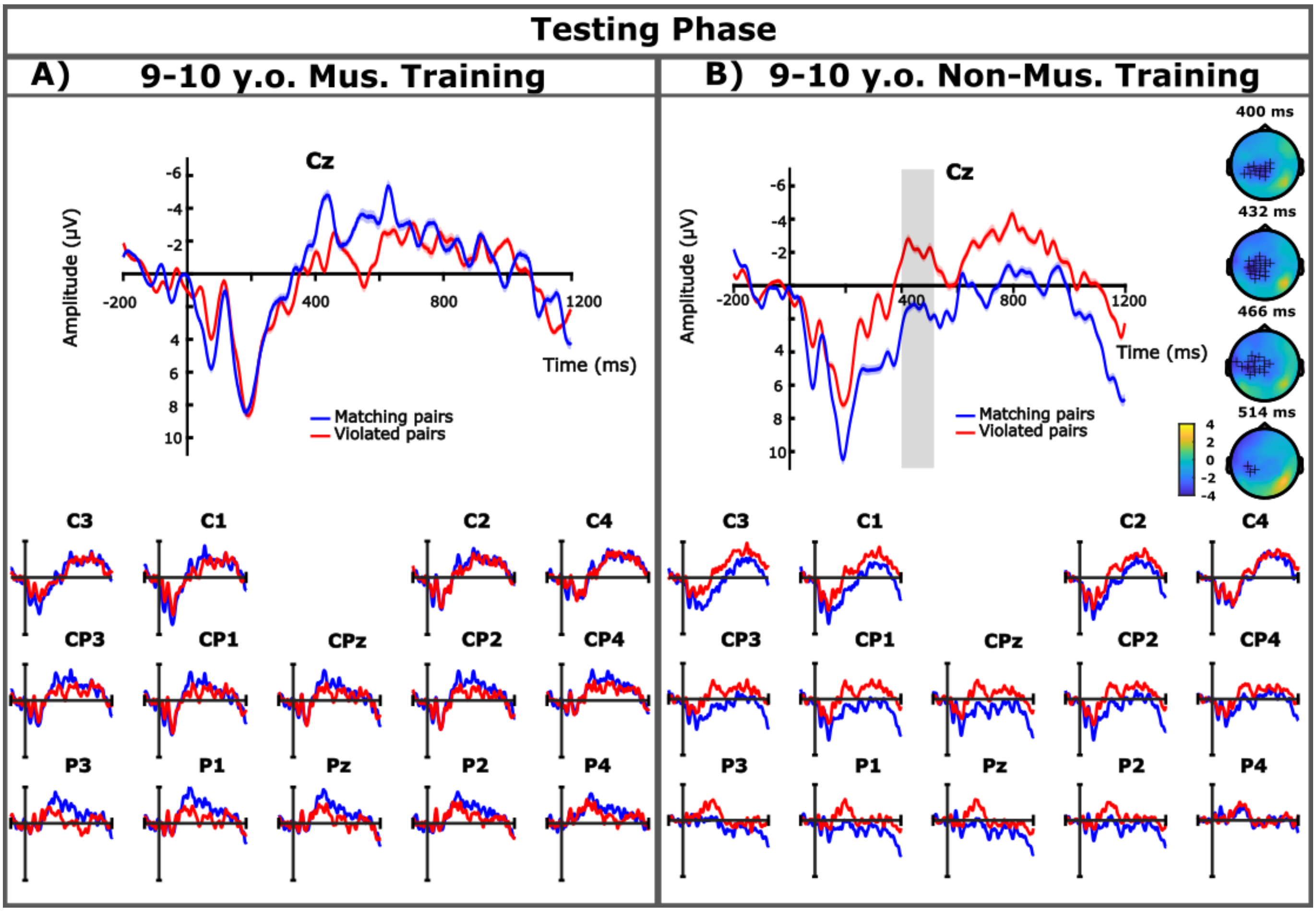


Figure SM_I_3: Testing phase results. ERPs are presented time-locked to the onset of the pseudoword for the testing phase of musically-trained and non-musically-trained 9-10 year olds. The Cz electrode was selected as a representative electrode for both groups, further centro-parietal electrodes are displayed below. The blue lines indicate the matching pairs while the red lines indicate the violated pairs. Gray bar indicates time window in which trend-level differences between the conditions were found. Topological difference maps from the cluster-based permutation test are presented only where trends-level differences were found and show the electrode distribution in the cluster over time (the + symbol indicates 0.1 < p < 0.05).

#### Musical-training differences: 9-10-year-olds testing phase results

For the testing phase, no clusters of significant differences for the musically trained participants were found in either time window (see Figure SM_I_3(A)). For the non-musically trained participants, while no clusters with significant differences was found in either time window, a negative trend was found in the later time window between 400 – 516 ms ($p$ = 0.092) (see Figure SM_I_3(B). While not significant, presentation of trend-level results is in line with previous publications between different groups in auditory associative word learning (cf. Experiment 2, Cosper et al., 2024).

#### J: Changes in AIC and model comparisons for the LMM of the behavioral data:

| *npar* | *AIC* | *BIC* | *logLik* | *deviance* | *Chisq* | *Df* | *Pr..Chisq.* |
|---|---|---|---|---|---|---|---|
| 3 | 193.99 | 205.74 | -93.99 | 187.99 | NA | NA | NA |
| 11 | -57.03 | -13.93 | 39.52 | -79.03 | 267.02 | 8 | 0.00 |

Table SM_J_1: Changes in AIC and BIC between the null model (top row) and the fit model (bottom row)

**Model comparison**

| | **Fit.model** | | **Null.model** | |
|---|---|---|---|---|
| *Predictors* | *Estimates* | *p* | *Estimates* | *p* |
| (Intercept) | 0.05 | 0.609 | 0.30 | **<0.001** |
| corrected hit-rate type [corrected_hit-rate_aud_nv] | -0.33 | **<0.001** | | |
| corrected hit-rate type [corrected_hit-rate_aud_ver] | -0.53 | **<0.001** | | |
| corrected hit-rate type [corrected_hit-rate_vis_nv] | -0.34 | **<0.001** | | |
| group | 0.09 | 0.418 | | |
| age | -0.02 | 0.533 | | |
| sex [m] | 0.05 | 0.100 | | |
| music | 0.02 | 0.574 | | |
| SETscore | 0.00 | **0.001** | | |
| **Random Effects** | | | | |
| $\sigma^2$ | 0.04 | | 0.09 | |
| $\tau_{00}$ | 0.01 subject | | 0.01 subject | |
| ICC | 0.21 | | 0.11 | |
| N | 93 subject | | 93 subject | |
| Observations | 372 | | 372 | |
| Marginal $R^2$ / Conditional $R^2$ | 0.488 / 0.596 | | 0.000 / 0.106 | |

Table SM_J_2: Model Comparisons between the null model and the fit model for the LMM of behavioral measures.

### K. Correlation Matrix for behavioral measures in table form:

*Means, standard deviations, and correlations with confidence intervals*

| Variable | *M* | *SD* | 1 | 2 | 3 | 4 | 5 | 6 | 7 | 8 | 9 | 10 | 11 | 12 | 13 | 14 | 15 |
|---|---|---|---|---|---|---|---|---|---|---|---|---|---|---|---|---|---|
| 1. group | 0.57 | 0.26 | | | | | | | | | | | | | | | |
| 2. age | 0.59 | 0.27 | .99** [.98, 1.00] | | | | | | | | | | | | | | |
| 3. sex | 0.03 | 0.27 | -.55 [-.82, -.07] | -.58* [-.84, -.12] | | | | | | | | | | | | | |
| 4. music | 0.25 | 0.24 | .24 [-.29, .66] | .24 [-.29, .66] | -.32 [-.70, .21] | | | | | | | | | | | | |
| 5. area1 | 0.53 | 0.25 | .86** [.63, .95] | .88** [.69, .96] | -.61* [-.85, -.17] | .16 [-.36, .61] | | | | | | | | | | | |
| 6. area2 | 0.48 | 0.27 | .76** [.43, .91] | .81** [.52, .93] | -.68** [-.88, -.28] | .03 [-.48, .51] | .88** [.69, .96] | | | | | | | | | | |
| 7. area3 | 0.50 | 0.27 | .89** [.70, .96] | .90** [.74, .97] | -.58* [-.84, -.12] | .09 [-.42, .56] | .80** [.49, .93] | .78** [.45, .92] | | | | | | | | | |
| 8. area4 | 0.57 | 0.27 | .91** [.75, .97] | .93** [.80, .98] | -.63** [-.86, -.19] | .03 [-.47, .52] | .87** [.65, .95] | .89** [.70, .96] | .89** [.71, .96] | | | | | | | | |
| 9. area5 | 0.42 | 0.27 | .80** [.51, .93] | .81** [.53, .93] | -.63** [-.86, -.19] | .01 [-.49, .50] | .76** [.43, .91] | .75** [.41, .91] | .70** [.32, .89] | .83** [.57, .94] | | | | | | | |
| 10. area6 | 0.42 | 0.23 | .57* [.11, .83] | .63** [.19, .86] | -.61* [-.85, -.16] | .23 [-.30, .65] | .72** [.34, .89] | .64** [.22, .86] | .55 [.07, .82] | .66** [.24, .87] | .53 [.05, .81] | | | | | | |
| 11. area7 | 0.34 | 0.23 | .37 [-.15, .73] | .39 [-.13, .74] | -.57* [-.83, -.11] | .12 [-.40, .58] | .47 [-.04, .78] | .48 [-.02, .79] | .35 [-.18, .72] | .34 [-.18, .72] | .35 [-.18, .72] | .38 [-.14, .74] | | | | | |

| | | | | | | | | | | | | | | | | | |
|---|---|---|---|---|---|---|---|---|---|---|---|---|---|---|---|---|---|
| 12. SETscore | 0.61 | 0.29 | .91** [.76, .97] | .94** [.84, .98] | -.70** [-.89, -.32] | .10 [-.42, .56] | .92** [.79, .97] | .92** [.78, .97] | .93** [.81, .98] | .96** [.89, .99] | .84** [.58, .94] | .70** [.32, .89] | .50 [.00, .80] | | | | |
| 13. corrected_hit rate_aud_ver | 0.26 | 0.24 | .14 [-.38, .59] | .11 [-.41, .58] | -.31 [-.70, .22] | -.58* [-.84, -.12] | .14 [-.38, .59] | .22 [-.30, .65] | .23 [-.30, .65] | .29 [-.24, .69] | .20 [-.33, .63] | .00 [-.50, .50] | .07 [-.44, .55] | .22 [-.31, .65] | | | |
| 14. corrected_hit rate_aud_nv | 0.45 | 0.20 | .54 [.06, .82] | .53 [.05, .81] | -.40 [-.75, .12] | .21 [-.32, .64] | .44 [-.07, .77] | .36 [-.17, .73] | .50 [.00, .80] | .56 [.08, .83] | .41 [-.11, .75] | .42 [-.09, .76] | .35 [-.18, .72] | .52 [.03, .81] | .10 [-.42, .57] | | |
| 15. corrected_hit rate_vis_nv | 0.41 | 0.21 | .27 [-.26, .68] | .30 [-.23, .69] | -.12 [-.58, .40] | .06 [-.45, .54] | .23 [-.30, .65] | .21 [-.32, .64] | .36 [-.17, .72] | .38 [-.15, .73] | .06 [-.45, .54] | .24 [-.29, .66] | -.10 [-.57, .42] | .28 [-.25, .68] | .05 [-.46, .53] | .53 [.05, .81] | |
| 16. corrected_hit rate_beh | 0.39 | 0.21 | .17 [-.35, .62] | .20 [-.33, .63] | -.14 [-.59, .39] | .08 [-.43, .55] | .21 [-.32, .64] | .19 [-.34, .63] | .26 [-.27, .67] | .31 [-.22, .70] | -.01 [-.50, .49] | .19 [-.33, .63] | -.12 [-.58, .40] | .21 [-.32, .64] | .04 [-.46, .53] | .43 [-.09, .76] | .82** [.54, .93] |

Table SM_K_1: Behavioral measures correlation matrix. *M* and *SD* are used to represent mean and standard deviation, respectively. Values in square brackets indicate the 95% confidence interval for each correlation. The confidence interval is a plausible range of population correlations that could have caused the sample correlation (Cumming, 2014). * indicates $p < .05$. ** indicates $p < .01$. FDR corrections have been applied to all *p*-values.
